# A method to disentangle enhancer-gene interactions from latent factors in single-cell multi-omics data

**DOI:** 10.64898/2026.11.24.768923

**Authors:** Ngoc Duc Nguyen, Michelle T. McNulty, Gyuhee Kim, Seong Kyu Han, Matthew G. Sampson, Dongwon Lee

## Abstract

Mapping regulatory elements to their target genes in the cell types relevant to disease is essential for interpreting non-coding variants identified by genome-wide association studies. Single-cell multiome data, which simultaneously measure accessible chromatin and gene expression per cell, allow such peak–gene links to be inferred within individual cell types. However, shared technical and biological variation across genes and peaks, such as cell cycle, cell quality, and inter-sample differences, creates spurious links, while including sample covariates to control for this variation inadvertently removes true genotype-driven links. To address this trade-off, we developed CREMAP (<u>C</u>ovariance-of-<u>R</u>esiduals-based <u>E</u>nhancer-gene <u>Map</u>ping), a statistical method that estimates shared variation as latent factors from the data and adjusts for them before testing each peak–gene pair. Using CRISPRi perturbation experiments and fine-mapped eQTLs as validation benchmarks, we demonstrate that CREMAP improves the accuracy of inferred peak–gene links compared with existing methods across two cell lines and 13 cell types from human bone marrow, brain, and kidney. In kidney proximal tubule cells, CREMAP retains eQTL- supported and allele-specific links that standard approaches miss when samples are included as covariates. Furthermore, its latent factors capture previously unrecognized global signatures, including a tubular injury program. CREMAP thus enables accurate cell-type-specific peak–gene mapping from single-cell multiome data while preserving genuine genetic regulatory effects.

## Introduction

Genome-wide association studies (GWAS) have identified thousands of genetic loci associated with complex traits and diseases^1,2^. Approximately 90% of the associated variants lie outside protein-coding sequences, predominantly within *cis*-regulatory elements such as enhancers and promoters^3^. These variants are thought to modulate regulatory activity and thereby alter target-gene expression in trait-relevant cell types^3,4^. For most loci, however, the underlying mechanism remains unresolved^5^. Two questions are central: the identity of the target genes, which are frequently not the nearest ones^6,7^, and the cell types in which regulation occurs, as genetic effects on expression often vary across cell types and cellular states^8,9^. Traditionally, expression quantitative trait loci (eQTLs) have been used to address these questions by linking genetic variants to changes in gene expression^10,11^. However, regulatory effects confined to specific cell types or cellular states often go undetected^12^ because bulk profiling averages signals across heterogeneous cell populations. Consistent with this, eQTLs mapped in bulk tissues account for only a minority of GWAS associations^4^.

Recently developed single-cell multiome technologies, which jointly profile gene expression and chromatin accessibility in individual cells, make it possible to address both questions directly^13,14^. Resolving constituent cell types from heterogeneous tissues allows peak–gene links to be inferred within each cell type by testing whether the accessibility of a regulatory element, measured as a peak in ATAC-seq (assay for transposase-accessible chromatin using sequencing) data, associates with target-gene expression across cells^15,16^. Moreover, multiome datasets are now generated at population scale, with cohorts reaching 10 million peripheral blood immune cells from 1,108 individuals^17^, allowing these cell-type-specific regulatory links to be analyzed alongside donor genotypes.

Computational methods that exploit this association fall into two classes: statistical linking methods and supervised integrative models. First, statistical linking methods infer peak–gene links from paired RNA-seq and ATAC-seq measurements alone by modeling the association between a peak’s accessibility and a gene’s expression across cells. They differ in how the relationship between the two modalities is quantified: by correlation across individual cells (Signac^18^; FigR^19^) or across aggregates of similar cells (ArchR^20^); by regression of one modality on the other (SnapATAC^21^; SCENT^22^; Open4Gene^23^; fasthurdle^17^); or by estimating their covariance under a count model that treats both modalities symmetrically (scMultiMap^24^). Second, supervised integrative models combine these linking scores with genomic distance and other features by learning feature weights from orthogonal validation data on enhancer–gene regulation. For example, pgBoost^25^, a gradient-boosting model trained on fine-mapped eQTLs, integrates the scores of SCENT, Signac, and Cicero^26^ with distance-based features. Similarly, scE2G^27^, a logistic-regression classifier trained on CRISPR perturbation data from K562 cells, combines an accessibility–expression correlation with the activity-by-contact score^28^ and other accessibility- and distance-based features.

All these approaches rely on this association across cells, but even within a cell type, genes and peaks also vary together for reasons unrelated to direct *cis*-regulation. Even in a single sample, cell-level factors such as cell-cycle stage, cell quality, and cell state simultaneously alter the expression of many genes and the accessibility of many peaks^29,30^. Moreover, in multi-sample studies, technical batch effects and biological differences across individuals introduce additional sample-level variation^31,32^. We refer to variation that is common to many genes and peaks across the cells of a cell type as shared variation. When it affects both modalities, genes and peaks covary regardless of whether they are linked by *cis*-regulation, producing spurious links. In multi-sample data, a standard remedy is to include sample identity as a covariate^22,24^. Adjusting for sample identity, however, removes all between-sample covariation, including covariation driven by *cis*-acting genetic variants that modulate peak accessibility and its target gene expression across donors^33^.

A similar problem arose in bulk expression studies, in which hidden factors such as batch, cell composition and sample quality affect thousands of genes. Estimating these factors from the data and removing their effects increases the power of eQTL mapping (surrogate variable analysis^34^; PEER^35^) and reduces false-positive edges in co-expression networks^36^, because such factors are shared by many features, whereas an individual regulatory link, or the effect of a *cis*-acting variant, involves only a few features. Here we apply this principle to peak–gene linking with **CREMAP** (<u>C</u>ovariance-of-<u>R</u>esiduals-based <u>E</u>nhancer-gene <u>Map</u>ping), which estimates latent factors that capture shared variation across the genome and adjusts for them in both modalities before testing each candidate pair. We benchmarked CREMAP against existing methods across two cell lines and 13 cell types from human bone marrow, brain, and kidney using CRISPRi perturbation experiments and fine-mapped eQTLs. In addition, we compared latent-factor adjustment with sample-covariate adjustment to evaluate their impact on genotype-driven links and characterized the technical and biological signals captured by the inferred latent factors.

## Results

### CREMAP removes shared variation and improves the accuracy of peak–gene links in cell lines

Our work was motivated by the observation that *trans* pairs, in which a gene and a peak reside on different chromosomes, are significantly correlated far more frequently than expected by chance. Because *trans* pairs cannot be linked in *cis*, only 5% would be expected to correlate at P < 0.05 under the null hypothesis. Instead, the fraction of correlated *trans* pairs exceeded this expectation in all 15 cell types across five datasets (5.7% to 22.7% of 20,000 randomly sampled pairs; **Fig. S1a**). This inflation was more prominent in cell types with more cells (ρ = 0.66; **Fig. S1b**). Through factor decomposition analysis, we found that this shared variation was low- dimensional: in each cell type, the peak–gene correlations had a few leading components, shared across many genes and peaks (**Fig. S1c**).

Inspired by this observation, we developed CREMAP to account for these global latent factors. CREMAP is a moment-based test of the covariance between a gene’s expression and a peak’s accessibility across the cells of a cell type. Building on the scMultiMap framework^24^, CREMAP estimates latent factors of shared variation from the cross-modality residual covariance across the genome and accounts for them before testing individual candidate pairs. Specifically, CREMAP works in four steps (**Fig. 1**): (1) a count model estimates the fitted mean and variance for each gene and peak given the cell’s sequencing depth, yielding residuals from these fits; (2) latent factors of shared variation are estimated from the covariance between gene and peak residuals across the genome; (3) the estimated latent factors are regressed out of both the gene and peak residuals; and (4) the covariance of each candidate pair is estimated and tested using the adjusted residuals. We first tested whether removing these latent factors reduces false positive correlations among *trans* pairs in two cell lines, K562 and GM12878. Regressing out the first two factors lowered the fraction of correlated *trans* pairs from 13.4% to 10.5% in K562 and from 13.2% to 7.6% in GM12878, removing a third and two-thirds of the excess over 5%, respectively (**Fig. 2a**).

**Figure 1.**
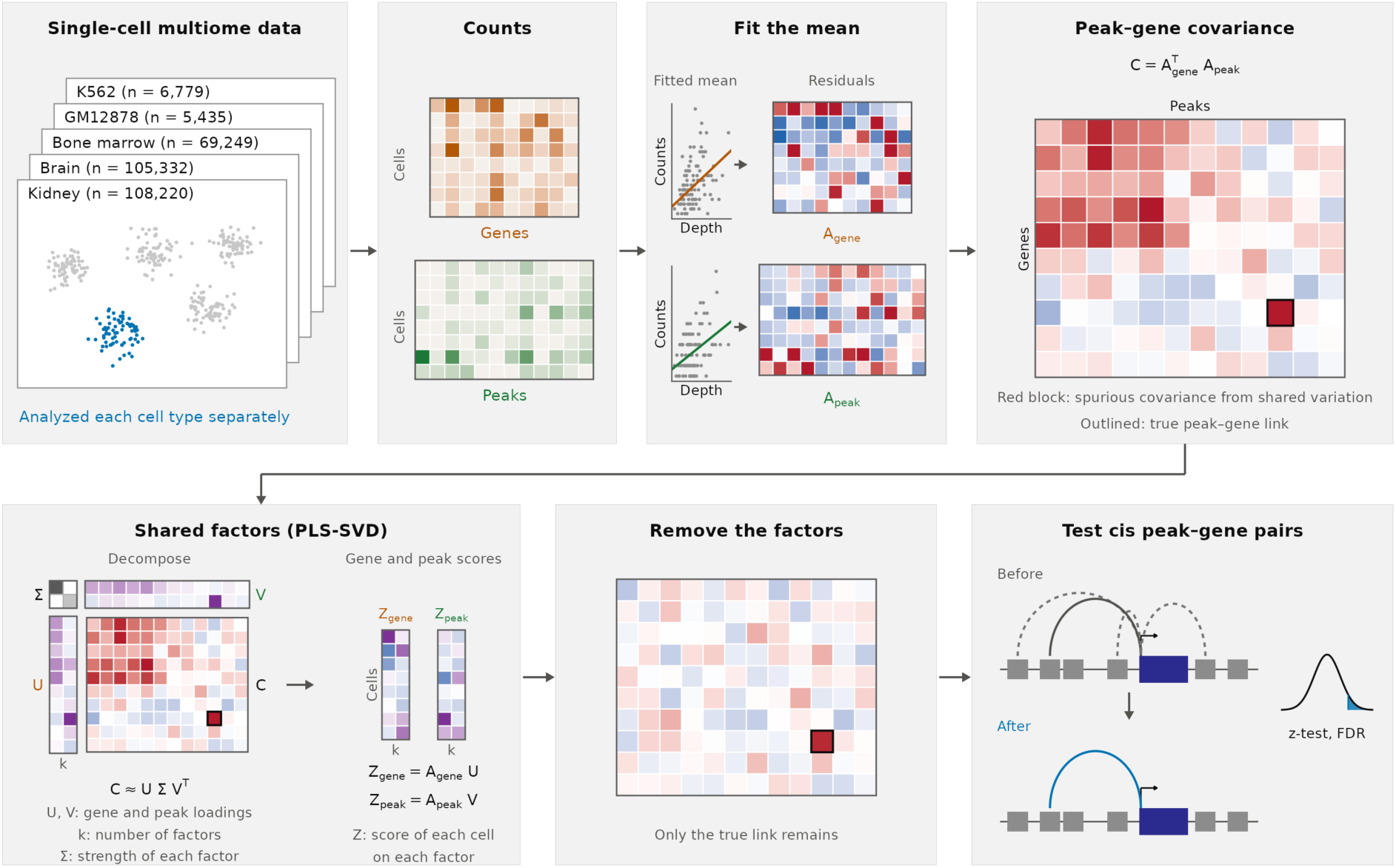
Schematic overview of CREMAP framework for predicting peak–gene links from single-cell multiome data. Within each cell type, CREMAP first fits the expected count given the cell’s sequencing depth to obtain gene and peak residual matrices, and then estimates shared variation across multiple peak–gene pairs as latent factors via partial least squares (PLS-SVD) to yield per-cell gene and peak scores. These scores are then regressed out of the residuals. Lastly, candidate peak–gene pairs are tested on the adjusted residuals using a z-test.

**Figure 2.**
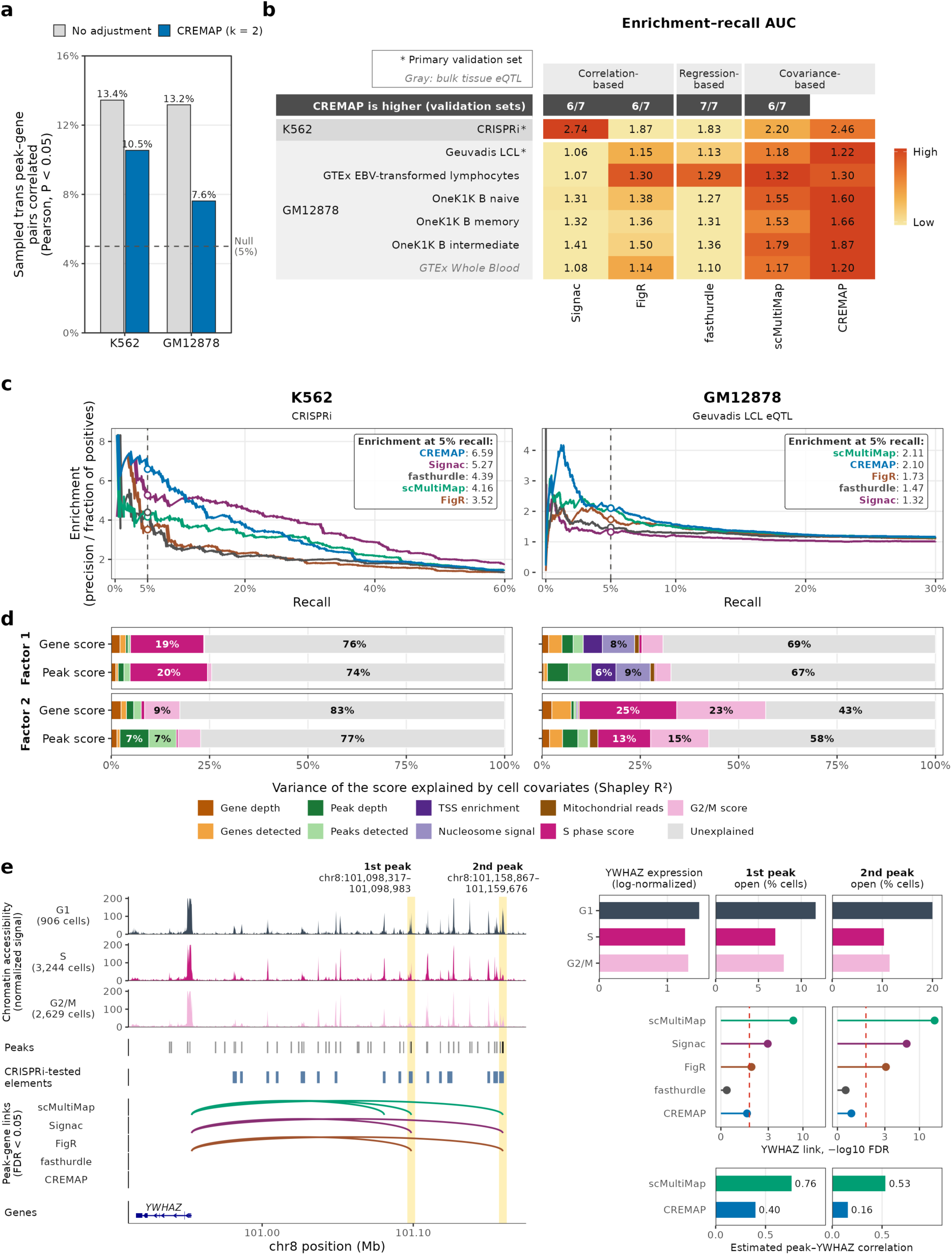
Performance of CREMAP in predicting peak–gene links in single-sample cell line data. **(a)** Percentage of 20,000 randomly sampled *trans* peak–gene pairs with significant Pearson correlation (P < 0.05) without adjustment (gray) and after adjusting for CREMAP’s latent factors (blue). The dashed line indicates the 5% null expectation. **(b)** Enrichment–recall AUC of five methods on K562 CRISPRi-tested element–gene pairs and six GM12878 eQTL sets. An asterisk (*) indicates a primary set for validation, used in (c). The color in the heatmap is relative to the best method in each row. The top row indicates the number of sets in which CREMAP’s AUC is higher than that of the method in the column. **(c)** Enrichment–recall curves for the primary validation sets in K562 (left) and GM12878 (right). The dashed line indicates 5% recall. **(d)** Variance of each latent factor’s gene and peak scores across cells explained by cell covariates (Shapley R²) in K562 (left) and GM12878 (right). Gray indicates unexplained variance, and shares of 6% or more are displayed. **(e)** *YWHAZ* locus in K562, where CRISPRi found no element regulating *YWHAZ* (65 elements tested within 1 Mb). The left panel shows, from top to bottom, the chromatin accessibility of G1, S and G2/M cells, peaks, CRISPRi-tested elements, each method’s peak–gene links (FDR < 0.05) and genes. Shaded bands mark the two focal peaks. The right panel shows *YWHAZ* expression and the percentage of cells with each peak open by cell cycle phase, each method’s FDR for the link of each peak (red dashed line, 0.05), and the correlation estimated by scMultiMap and CREMAP.

We next asked whether accounting for these latent factors improves peak–gene linking. We compared CREMAP with four other existing methods (**Methods**): Signac and FigR correlate accessibility with expression and compare each pair with background peaks, which accounts for peak accessibility and GC content but not for variation shared across the genome; fasthurdle regresses expression on accessibility using a two-component hurdle model and adjusts only for sequencing depth and user-supplied covariates; and scMultiMap estimates covariance between peak accessibility and gene expression in a similar manner to CREMAP. To evaluate the performance, we calculated the area under the enrichment–recall curve (enrichment–recall AUC), where enrichment is defined as precision relative to random chance (the ratio of validated links among predicted pairs to the baseline validation rate across all candidate pairs) (**Methods**). Using CRISPRi-validated pairs in K562 and six eQTL sets in GM12878 as benchmarks, we found that CREMAP achieved a higher enrichment–recall AUC than scMultiMap, Signac and FigR in 6 of the 7 validation sets, and outperformed fasthurdle in all 7 (**Fig. 2b**; **Table S1**). At 5% recall, CREMAP achieved 6.6-fold enrichment compared to 5.3-fold for the second-best method (Signac) in K562, and matched scMultiMap (2.10–2.11) while exceeding the third best method (FigR, 1.73) in GM12878 (**Fig. 2c**). We also examined which subsets of peak–gene links benefited most from latent-factor adjustment. We stratified candidate pairs by gene expression, peak accessibility signal, and distance to TSS and found that the performance gain was most consistent for highly expressed genes and distal (>100 kb) peak–gene pairs (**Fig. S2a–d**). Finally, among the links predicted by the other methods, CREMAP preferentially retained validated links over non-validated ones. Similarly, among the candidate pairs that Signac, FigR and fasthurdle missed, CREMAP still recovered validated pairs more often than non-validated ones (**Fig. S2e, f**).

### CREMAP identifies technical and biological latent factors

We next asked what types of signals are captured by these latent factors. Because CREMAP estimates each factor jointly from cross-modality residual covariance, projecting individual cells onto each factor’s gene and peak loadings yields two modality-specific cell scores per factor, which we refer to as gene and peak factor scores (**Methods**). We partitioned the variance of each factor’s gene and peak scores among per-cell measurements and found that the cell-cycle scores explained the largest proportion of variance in K562’s first factor and GM12878’s second factor (19–48%; **Fig. 2d**; **Fig. S3a**). In both factors, the genes with the highest loadings were enriched for cell-cycle gene sets, such as E2F targets and G2/M checkpoint (**Fig. S3b, c**; **Table S2**). Because cell-cycle and other global programs simultaneously alter chromatin accessibility and gene expression, they can induce spurious correlations between unlinked peaks and genes. By accounting for them, these spurious links can be effectively controlled, even if they are in *cis*. As a representative example, we evaluated candidate pairs at the *YWHAZ* locus in K562, where CRISPRi perturbation found no regulatory elements among 65 tested peaks targeting *YWHAZ*. At this locus, scMultiMap, Signac, and FigR linked the gene to two distal peaks whose accessibility correlated with cell cycle phase, which in turn correlated with *YWHAZ* expression. In contrast, CREMAP accounted for this cell-cycle derived signal and assigned non-significant P values to both candidate pairs (**Fig. 2e**). We observed the same pattern at two additional loci. In K562, scMultiMap linked a single peak to three genes, none of which were validated as targets by CRISPRi, whereas CREMAP did not call any of these links (**Fig. S4a**). Similarly, at *SERPINB9* locus in GM12878, all four existing methods linked the gene to a peak whose accessibility correlated with a global TNF-α/NF-κB program, while CREMAP removed that spurious link and kept the link to the peak harboring the gene’s fine-mapped eSNP and promoter (**Fig. S4b**). Together, these results demonstrate that CREMAP successfully identifies global latent factors and, by accounting for them, it reduces false-positive peak–gene links driven by shared cellular programs.

### Using the sample as a covariate removes eQTL-supported and genotype-driven peak–gene links

We next evaluated the impact of including sample identity as a covariate in existing peak–gene linking methods, as recommended by tools such as scMultiMap and SCENT, to control for sample-level confounding factors, particularly in multi-sample cohorts. Although this adjustment is well motivated for removing technical batch effects and broad biological differences across individuals, we reasoned that regressing out sample identity also eliminates genuine *cis*-genetic variation across donors, thereby discarding true peak–gene links supported by eQTLs.

To test this, we ran scMultiMap and fasthurdle both with and without a sample covariate in 13 cell types from three multi-sample datasets (5 bone-marrow, 6 brain and 2 kidney cell types) and evaluated the resulting peak–gene link maps against eQTL datasets matched to the corresponding cell type (**Methods**). Consistent with our expectation, adding the sample covariate lowered recall at FDR < 0.05 in all 13 cell types for both methods, dropping from a median of 6.4% to 1.6% for scMultiMap and from 2.4% to 0.2% for fasthurdle. Overall ranking accuracy also declined, with the enrichment–recall AUC decreasing in 12 of the 13 cell types for scMultiMap and in 11 for fasthurdle (**Fig. 3a**). To examine this loss in greater detail, we focused on kidney proximal tubule (PT) cells. In PT cells, adding the sample covariate eliminated 79% of the links originally detected by scMultiMap (**Fig. 3b**), including 66% of those supported by fine-mapped eSNPs from the NEPTUNE TUBE eQTL dataset (**Fig. 3c**)^37^. Even after controlling for distance to the TSS, sequencing depth, and peak width, lost links showed nearly identical enrichment for eQTL support as retained links (odds ratio versus non-linked pairs, 2.44 and 2.39, respectively; **Fig. 3d**), indicating that they were not merely technical or genomic artifacts. We observed the same enrichment for lost links when evaluating another independent cohort, GTEx Kidney Cortex (**Fig. S5a**).

**Figure 3.**
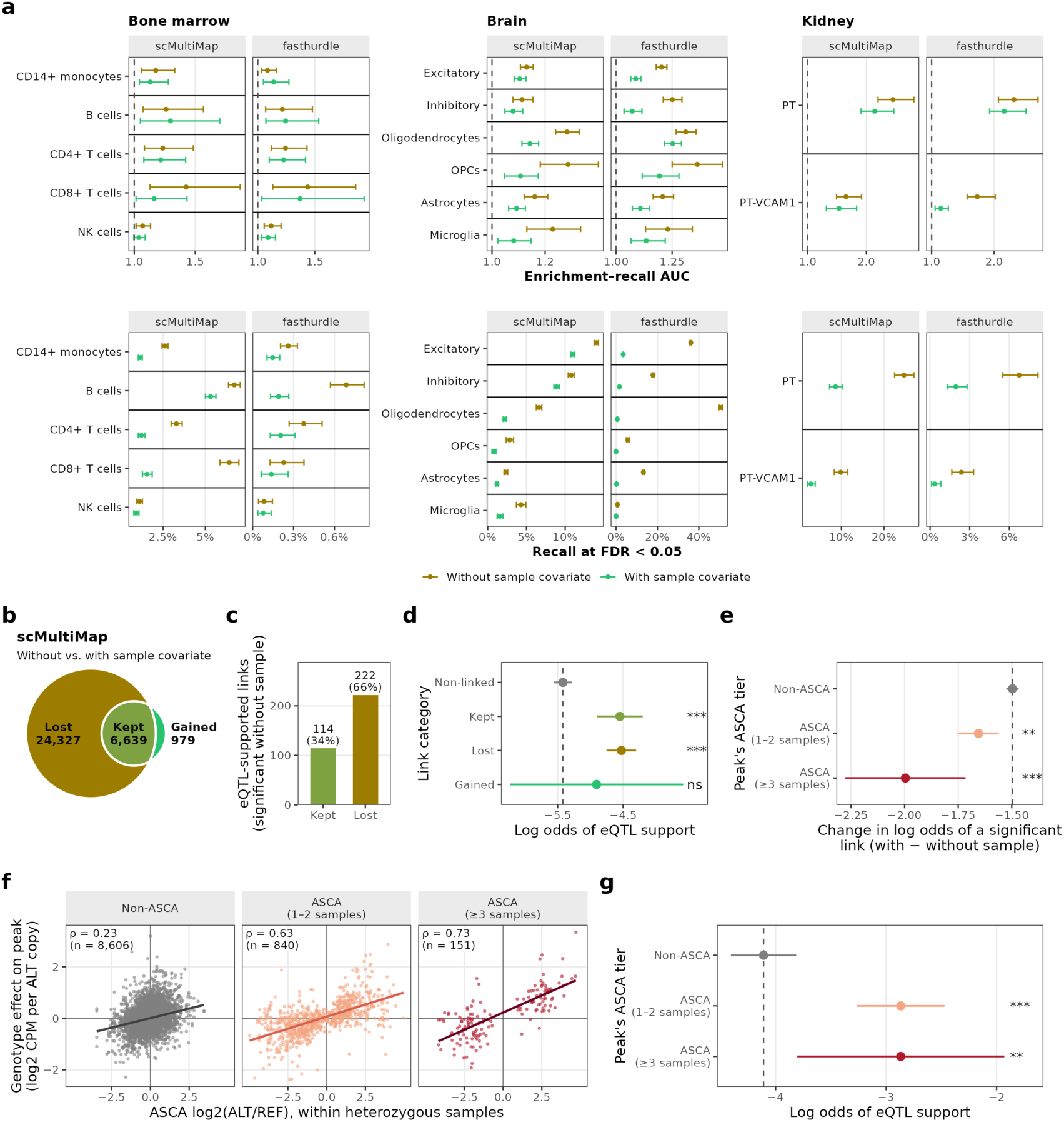
The effect of using sample identity as a covariate in peak–gene linking models. **(a)** Enrichment–recall AUC (top) and recall at FDR < 0.05 (bottom) of cell-type-matched eQTLs for scMultiMap and fasthurdle without (mustard) and with (green) the biological sample as a covariate, in 13 cell types from three multi-sample datasets (whiskers, 95% CI; dashed line, AUC = 1, indicating no enrichment). Panels (b)–(g) show scMultiMap in kidney proximal tubule (PT) cells. **(b)** Links significant (FDR < 0.05) without the sample covariate only (Lost), in both models (Kept) or with the sample covariate only (Gained). **(c)** Kept and lost links supported by fine-mapped NEPTUNE TUBE eSNPs. **(d)** Log odds that a pair is eQTL-supported by link category (Non-linked, significant in neither model), after adjusting for distance to the TSS, sequencing depth and peak width. **(e)** Change in the log odds that a candidate pair is a significant link when the covariate is added, by the peak’s allele-specific chromatin accessibility (ASCA) tier, adjusted for allelic coverage and sequencing depth. **(f)** For peaks that lost a link, ASCA log2(ALT/REF) within heterozygous samples against the genotype effect on accessibility across 22 samples. Lines show least-squares fits, ρ indicates Spearman correlation, and n indicates the number of peaks. **(g)** Log odds that a lost link is eQTL-supported by ASCA tier (adjusted as in d). In (d), (e) and (g), dots and whiskers show model estimates with 95% CI, dashed lines indicate the reference category (d, Non-linked; e and g, Non-ASCA), and asterisks indicate P versus the reference (* P < 0.05; ** P < 0.01; *** P < 0.001; ns, not significant).

To further test whether genotype explains these lost links directly, we identified variants with allele-specific chromatin accessibility (ASCA) and asked whether these variants are enriched in these lost links. Because ASCA compares chromatin accessibility between the two alleles at a heterozygous site within the same sample, it is unaffected by between-sample differences and provides an orthogonal benchmark of genuine *cis*-genetic effects. We first performed genome-wide ASCA variant calling in PT cells (**Methods**). We then called a peak an ASCA peak if it harbored at least one significant ASCA variant in at least one sample, and a replicated-ASCA peak if that variant was significant in at least three samples with a consistent allelic direction. Adding the covariate lowered the odds of a link being significant more at ASCA peaks than at non-ASCA peaks, and most strongly at replicated-ASCA peaks (**Fig. 3e**). If these lost links reflected technical differences between samples, accessibility across samples would not follow the genotype at individual SNPs. Instead, at peaks that lost a link, accessibility followed genotype in the direction predicted by ASCA (Spearman ρ = 0.73 at replicated-ASCA peaks versus 0.23 at non-ASCA peaks; **Fig. 3f**), as did expression of the linked gene, in the direction implied by the link (**Fig. S5b–d**). Lost links at ASCA peaks also showed the highest rate of eQTL support in the orthogonal NEPTUNE TUBE dataset (**Fig. 3g**). Repeating these analyses with fasthurdle yielded consistent results (**Fig. S6a–f**). Thus, including sample covariates indiscriminately removes between-sample variation, discarding true regulatory peak–gene links driven by *cis*-genetic variation across individuals.

### CREMAP preserves genotype-driven links in multi-sample data

We next hypothesized that latent factors estimated from genome-wide shared variation in CREMAP would not capture genotype-driven links, because a *cis*-acting variant alters only its local peak and target gene, with genotypes varying independently across the genome. To test this, we ran CREMAP in the same 13 cell types and compared it with the sample-adjusted runs from **Fig. 3**. We also included Signac and FigR, which do not support sample covariates and were therefore run without adjustment. As expected, CREMAP achieved a higher enrichment–recall AUC than sample-adjusted scMultiMap in all 13 cell types and sample-adjusted fasthurdle in 11, while also achieving higher recall at FDR < 0.05 in 8 and 13 cell types (**Fig. 4a**). Moreover, compared with unadjusted Signac and FigR, CREMAP achieved higher recall in all 13 cell types and a higher enrichment–recall AUC in 11 and 7, respectively. The performance gain by CREMAP was the most prominent when calling distal peak–gene pairs, as peaks >20 kb from target genes’ TSS were more accurately called by CREMAP for the majority of the datasets compared to other methods (**Fig. S7a–d**). To evaluate the robustness of these enrichment results, we repeated the analyses across all 51 eQTL validation sets matched to these cell types (**Fig. S8**). CREMAP’s enrichment–recall AUC remained higher than that of sample-adjusted scMultiMap in 43 of the 51 sets and sample-adjusted fasthurdle in 33 sets. At FDR < 0.05, CREMAP’s recall was comparable to sample-adjusted scMultiMap and exceeded all three other methods, outperforming Signac, FigR, and sample-adjusted fasthurdle in 45, 46, and 51 of 51 validation sets, respectively.

**Figure 4.**
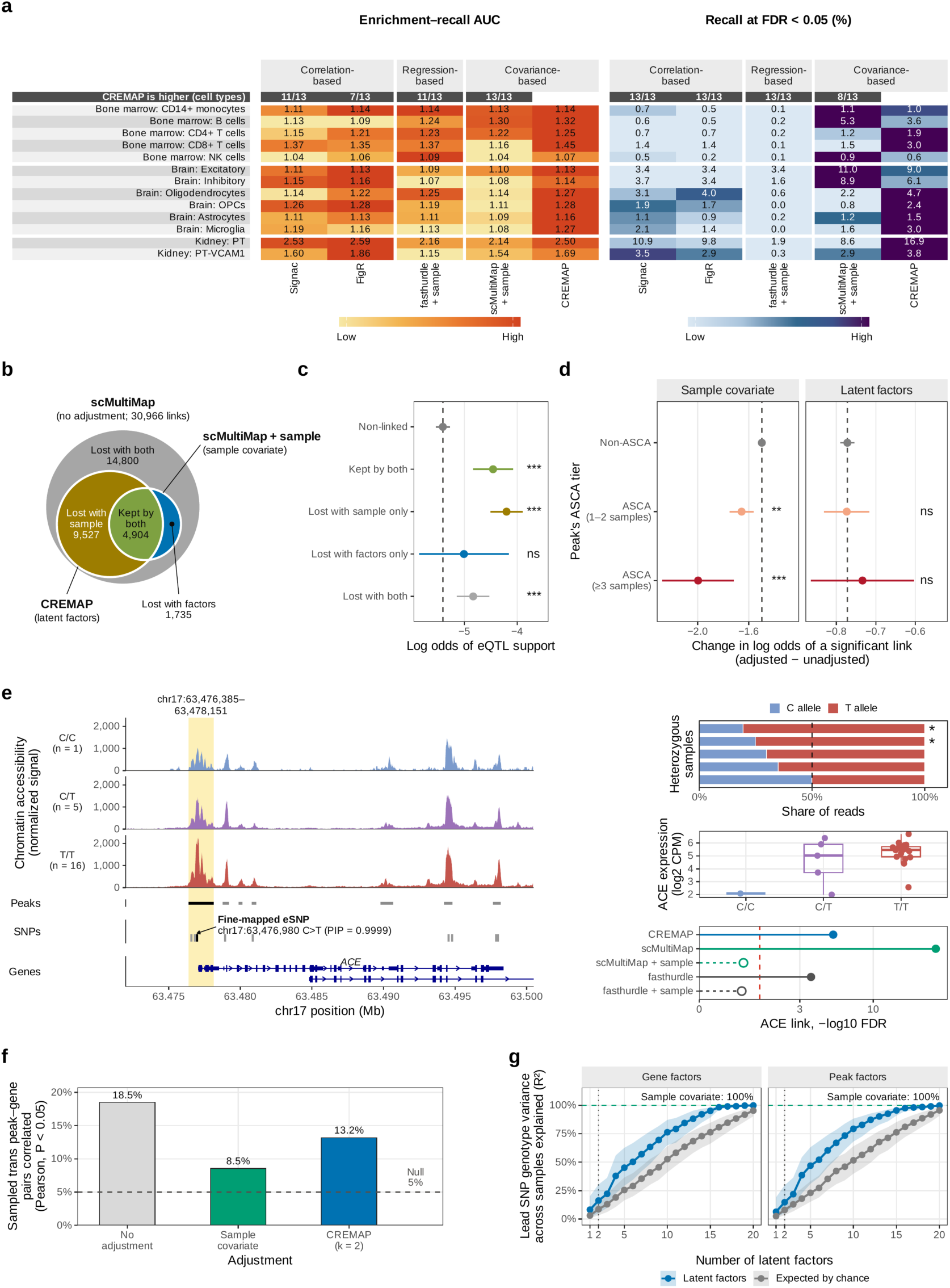
Performance of CREMAP in predicting peak–gene links in multi-sample data. **(a)** Enrichment–recall AUC (left) and recall at FDR < 0.05 (right) against each cell type’s primary eQTL validation set across 13 cell types. Color is relative to the best method in each cell type, and the top row shows the number of cell types where CREMAP achieves a higher value. Panels (b)–(g) show results in kidney proximal tubule (PT) cells. **(b)** Overlap of significant links (FDR < 0.05) called by unadjusted scMultiMap, scMultiMap with a sample covariate, and CREMAP. **(c)** Log odds of eQTL support by link fate under the two adjustments (dashed line, non-linked pairs). **(d)** Change in log odds of a significant link after each adjustment, by the peak’s ASCA tier (dashed line, Non-ASCA; x-axis scales differ between panels). In (c) and (d), dots and whiskers show estimates with 95% CI, and asterisks denote significance versus the reference category (* P < 0.05; ** P < 0.01; *** P < 0.001; ns, not significant). **(e)** *ACE* locus. Left: chromatin accessibility by rs4292 genotype, peaks (focal peak in yellow), tested SNPs including the fine-mapped eSNP, and genes. Right: allelic read share in heterozygous samples (* ASCA FDR < 0.05), *ACE* expression by genotype (boxes show median and quartiles), and link significance across methods (red dashed line, FDR = 0.05, with open dots and dashed lines indicating a sample covariate). **(f)** Percentage of 20,000 sampled *trans* peak–gene pairs significantly correlated across cells (Pearson P < 0.05) under each adjustment, compared with the 5% null expectation (dashed line). **(g)** Variance in lead SNP genotypes across 23 samples explained by the first *k* gene (left) or peak (right) latent factors, compared with permuted genotypes (median and quartiles across 23,317 lead SNPs).

We next examined how CREMAP’s latent factors and sample-covariate adjustment partition technical and biological variation in PT cells. Of the 30,966 links called by unadjusted scMultiMap, CREMAP’s latent factors removed 53%, whereas sample-adjusted scMultiMap removed most of these along with an additional 31% of links, leaving 5.6% of links removed exclusively by CREMAP’s latent factors (**Fig. 4b**). Crucially, links removed by the latent factors alone were not significantly enriched for eQTL support, in contrast to those removed exclusively by the sample covariate (**Fig. 4c**). Moreover, link loss under sample covariate adjustment was largest at replicated-ASCA peaks, whereas link loss under CREMAP’s latent factors was comparable between ASCA and non-ASCA peaks (**Fig. 4d**). Across all candidate pairs in PT cells, CREMAP’s links at FDR < 0.05 captured 17.5% of the 1,324 eQTL-supported pairs, compared with only 8.9% and 2.1% for sample-adjusted scMultiMap and fasthurdle (**Fig. S9a**; **Table S3**). While unadjusted Signac and FigR ranked these pairs comparably to or slightly higher than CREMAP among their top ∼10,500 and ∼13,500 predictions, they recovered fewer validated pairs at FDR < 0.05 (11.3% and 10.1%, respectively). Lastly, CREMAP’s links were enriched at replicated-ASCA peaks (odds ratio 1.57 against non-ASCA peaks), significantly more than those of sample-adjusted scMultiMap and fasthurdle (interaction P = 3 × 10^-4^ to 0.01; **Fig. S9b**; **Methods**).

A representative example illustrates the mechanism underlying such genotype-driven links. The *ACE* gene encodes angiotensin-converting enzyme, a prominent surface protein of the proximal tubule brush border^38^. Genotype variation at this locus explains nearly half of the variance in circulating ACE levels^39^, and therapeutic inhibition of the enzyme mitigates chronic kidney disease progression^40,41^. The fine-mapped eQTL for *ACE* in NEPTUNE TUBE (rs4292; posterior inclusion probability 0.9999) resides 81 bp upstream of the *ACE* TSS within the promoter peak and displays an ASCA effect: in four of five heterozygous samples, more ATAC reads carried the T allele. Across all samples, the T allele was associated with higher peak accessibility and *ACE* expression (**Fig. 4e**). Accordingly, this peak–gene link was significant in CREMAP (FDR = 5 × 10^-6^) as well as in unadjusted scMultiMap and fasthurdle, but lost after adding the sample covariate (sample- adjusted scMultiMap FDR = 0.15). At this locus, the between-sample variation removed by the sample covariate reflects an eQTL-driven regulatory effect, which CREMAP preserved.

### CREMAP removes shared variation between samples while preserving genetic variation

We next asked whether CREMAP, while keeping genotype-driven links, still removes shared variation in multi-sample data. We evaluated the correlation of *trans* pairs as in **Fig. 2a**. Without adjustment, 18.5% of 20,000 sampled *trans* pairs in PT cells were correlated (P < 0.05), whereas CREMAP’s latent factors reduced this inflation to 13.2% compared with 8.5% with a sample covariate (expected 5% under the null hypothesis; **Fig. 4f**). The sample covariate removed more of this inflation because it removes all between-sample variation, including genotype-driven variation. By construction, it accounts for 100% of the genotype variance across samples, whereas CREMAP’s two factors explained 15–16% of the genotype variance of lead SNPs across the 23 samples, close to the 8–9% expected by chance (**Fig. 4g**).

To identify what the latent factors captured, we decomposed the variance of gene and peak scores across known biological and technical covariates (**Fig. S10a**). Sample accounted for the largest proportion of variance in both factors (20–40% of score variance), yet the underlying drivers of these factors were distinct. The first factor’s gene score tracked RNA sequencing depth, with RNA counts and genes detected per cell explaining a further 36% of its variance; its mean per sample correlated with mean RNA sequencing depth (Pearson r = 0.58) and did not differ significantly between healthy (n = 18) and chronic kidney disease (CKD, n = 5) samples (P = 0.23). Conversely, the second factor showed no correlation with RNA sequencing depth (Pearson r = −0.13) but was elevated in CKD samples (P = 0.03). Gene set enrichment analyses agreed with this distinction: the first factor showed little enrichment for biological pathways (5 of 50 Hallmark sets at FDR < 0.05), consistent with a technical factor, whereas the second factor separated 13 injury and inflammation sets (including TNF-α signaling via NF-κB, inflammatory response and TGF-β signaling) from 10 sets of normal tubular metabolism (including oxidative phosphorylation and fatty acid metabolism), a pattern described in injured and diseased kidney tubules^42,43^ (**Fig. S10b**). Thus, CREMAP’s first factor captures RNA sequencing depth, while the second captures a tubular injury and inflammation program that was higher in CKD samples; both represent shared variation across many genes and peaks, which CREMAP removes without discarding genotype-driven variation.

## Discussion

Here we introduce CREMAP, a statistical method that removes the shared variation that inflates peak–gene associations in single-cell multiome data. CREMAP improved the accuracy and recall of peak–gene links across multiple cell types and tissues compared with existing methods, particularly for distal links. In multi-sample datasets, it also retained the eQTL-supported and allele-specific links that a conventional sample covariate removed. Finally, we demonstrated that decomposing the latent factors into per-cell measurements reveals which biological and technical properties confound peak–gene association.

Our work has direct implications for analytical strategies in multi-sample multiome studies. While incorporating a sample covariate was originally designed for differential expression across donors to account for the correlation of cells from the same donor^32^, existing peak–gene linking methods adopt this adjustment routinely into multi-sample datasets^22,24^. In peak–gene linking, however, differences between samples are not purely technical or global, as they also carry *cis*-genetic effects that a sample covariate removes together with non-genetic variation. Bulk eQTL mapping avoids this conflict by modeling genotype explicitly as the explanatory term, while removing global variation through hidden factors^34,35^. CREMAP applies a similar principle by separating shared variation common to many genes and peaks from locus-specific *cis*-genetic variation that links individual peaks to their target genes.

Beyond the sample, current linking methods either adjust for user-specified covariates, such as sequencing depth or cell quality (SCENT^22^; scMultiMap^24^; fasthurdle^17^), or test each pair against background peaks matched for GC content and accessibility (Signac^18^; FigR^19^). Both approaches require sources of variation to be anticipated in advance, and any unspecified confounders, such as the cell cycle or a cell state, remain uncorrected. CREMAP, by contrast, offers a framework that requires no prior specification and can discover previously unrecognized technical and biological sources of shared variation.

Our study has several limitations that point to future work. First, CREMAP is a count-based model and therefore depends on sequencing depth; methods that binarize chromatin accessibility^22,27^ may be more robust in sparse data, and applying CREMAP to other paired RNA-seq and ATAC-seq assays, such as SNARE-seq^13^ and SHARE-seq^14^, will test its generalizability. Second, removing two latent factors was not equally effective across cell types, indicating the need for a strategy to optimize the number of factors for each dataset. Third, our evaluation is limited by the validation data currently available to the field, as large-scale CRISPRi screens of enhancer–gene pairs exist mainly for K562^44^, and eQTLs indicate association rather than causality. Thus, experimental validation of CREMAP’s links in relevant cell types remains essential. Finally, whether incorporating CREMAP’s scores improves supervised integrative models such as pgBoost^25^ and scE2G^27^ remains to be tested. Addressing these points will further strengthen the use of single-cell multiome data for mapping non-coding variants to their target genes.

## Methods

### CREMAP

CREMAP extends scMultiMap^24^, a moment-based test of the covariance between a gene’s expression and a peak’s accessibility across the cells of a cell type. It keeps scMultiMap’s count model (equation (1)) and covariance test (equation (4)), and adds two steps: latent factors of shared variation are estimated from the covariance between gene and peak residuals across the genome (Step 2) and removed from the residuals (Step 3).

#### Count mode

In a cell type with *n* cells, let *x*_g*i*_ and *x_pi_* be the counts of gene *g* and peak *p* in cell *i*, and *s_i_* and *t_i_* the cell’s gene depth and peak depth (its total gene and peak counts). As in scMultiMap, the counts follow a Poisson distribution given the cell’s underlying expression and accessibility, so that

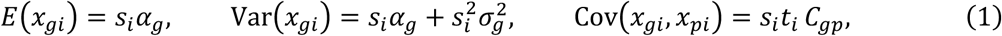

and likewise for peaks, with *t_i_*, *α_p_* and ‘[inlin]. Here *α*_g_ is the mean expression of gene *g* per unit depth, *σ*^2^ is the variance of its underlying expression across cells, and *C_gp_* is the covariance between the underlying expression of gene *g* and the underlying accessibility of peak *p*.

#### Shared variation

External factors, such as cell quality, cell-cycle phase, or between-sample differences add covariance to multiple gene–peak pairs, whether or not the pair is linked in *cis*. CREMAP estimates the shared variation from the data as a small number (*k*) of latent factors and removes this shared variation.

#### Step 1: residuals

For each gene, *α*_g_ and 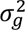 are estimated from the mean and variance conditions of (1) by weighted least-squares regressions on *s_i_* and 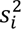, with negative-binomial variance weights *w*_g*i*_ updated by iteratively reweighted least squares; peaks are fitted in the same way with *t_i_*. The residuals are *r*_g*i*_ = *x*_g*i*_ − *s_i_α̂*_g_ and *r_pi_* = *x_pi_* − *t_i_α̂_p_*.

#### Step 2: latent factors

Let *A*_gene_ (*n* × *n*_gene_) and *A*_peak_ (*n* × *n*_peak_) denote the matrices of gene and peak residuals across the *n* cells, with each residual centered and scaled to unit variance, where *n*_gene_ and *n*_peak_ are the numbers of genes and peaks. The cross-modality matrix

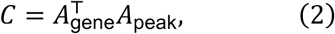

is then proportional to the correlation between every gene residual and every peak residual. Because almost all gene–peak pairs are not linked, the leading structure of *C* reflects shared variation. The first *k* pairs of singular vectors of *C* (partial-least-squares singular value decomposition; Tucker^45^) give gene loadings *U*_j_ and peak loadings *V*_j_ for each factor *j*, and each cell receives a gene score *Z*_gene,j_ = std(*A*_gene_*U*_j_* and a peak score *Z*_peak,j_ = std(*A*_peak_*V*_j_*, standardized to mean 0 and standard deviation 1. The gene score measures how strongly factor *j* is present in a cell’s gene expression residuals, and the peak score how strongly it is present in the cell’s chromatin accessibility residuals.

#### Step 3: removal of the factors

The factor scores of each modality are regressed out of that modality’s residuals by ordinary least squares:

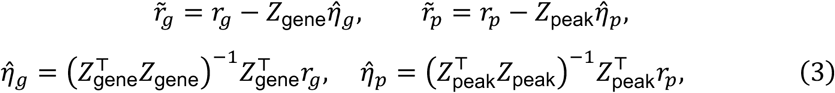

where *r*_g_ and *r_p_* are the residual vectors of gene *g* and peak *p* across cells, *Z*_gene_ and *Z*_peak_ are the *n* × *k* matrices of gene and peak scores, and *η̂_g_* and *η̂_p_* are the ordinary least-squares coefficients of the residuals on the scores.

#### Step 4: test

For a candidate pair (*g*, *p*), with *d_i_* = *s_i_t_i_* and *w_i_* = *w*_g*i*_*w_pi_*, *C*_g*p*_ is estimated by weighted least squares from the covariance condition of (1) applied to the adjusted residuals and tested with

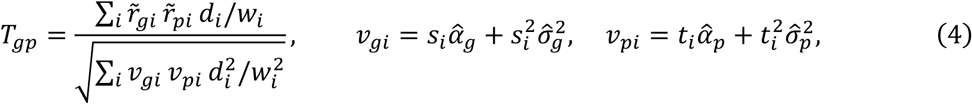

which is asymptotically standard normal under the null hypothesis of no association^24^.

### Datasets and preprocessing

For single-sample datasets, publicly available 10x Genomics Multiome (ATAC + Gene Expression) data from two cell lines, K562 and GM12878, were analyzed. For K562, raw FASTQ sequencing files of the 10x Multiome dataset were obtained from Xu et al.^46^ (ArrayExpress E-MTAB-11264) and were processed by Cell Ranger ARC 2.0.2 (GRCh38-2020-A). Cells meeting the following criteria were retained for the analysis (n=6,779): (1) ≥1,000 UMIs, (2) ≥ 2,000 genes, (3) ≤ 20% mitochondrial gene counts, (4) 1,000 ≤ ATAC-seq fragments ≤ 100,000 and (5) TSS enrichment ≥ 5. For GM12878, Cell Ranger ARC outputs of the 10x Multiome dataset from Wu et al.^47^ (GEO GSE166797) were obtained, and cells meeting the following criteria were retained (n=5,435): (1) 500 ≤ UMIs ≤ 50,000, (2) 200 ≤ gene ≤ 8,000, (3) ≤25% mitochondrial gene counts, (4) 1,000 ≤ peak counts ≤ 100,000, (5) TSS enrichment score ≥ 2, and (6) nucleosome signal ≤ 2.

For multi-sample datasets, datasets from three different human tissues (bone marrow, brain, and kidney) were examined. First, preprocessed and annotated cells from the NeurIPS 2021 bone marrow mononuclear cells multiome dataset^48^ (GEO GSE194122) were used directly without re-annotation (69,249 cells across 13 samples from 10 donors at 4 sites with sample identity used as the sample variable). Five major cell-type groups were defined from the original cell annotations for our analysis: B cells (naive CD20+, B1, transitional B, and plasma cells; 10,131 cells), CD14+ monocytes (10,843), CD4+ T cells (naive and activated; 9,924), CD8+ T cells (CD8+ T and naive CD8+ T; 12,601) and natural killer cells (6,929). Because fragment files were not included in the processed release, ATAC-seq fragments were regenerated from their aligned reads (ENA SRR17693253 to SRR17693265) using sinto 0.10.1. Second, for brain, we obtained the dorsolateral prefrontal cortex multiome dataset from Anderson et al.^49^ (GEO GSE214979), comprising processed and annotated 105,332 nuclei from 15 donors (7 with Alzheimer’s disease and 8 controls, analyzed together, with the donor as the sample variable). These nuclei were used without re-annotation across six cell types, including excitatory neurons (39,884 nuclei), inhibitory neurons (16,331), oligodendrocytes (34,784), oligodendrocyte precursor cells (OPCs; 4,193), astrocytes (6,255) and microglia (3,179). Third, for kidney, we analyzed 23 biological samples from 18 donors processed as part of a separate study (6 from Wilson et al.^50^, GEO GSE232222; 12 from Ledru et al.^51^, and 5 newly generated in our laboratory). Raw FASTQ files were uniformly processed with Cell Ranger ARC (GRCh38-2020-A), and cells meeting the following criteria were retained: (1) ≥ 1,000 ATAC-seq fragments, (2) TSS fragment fraction ≥ 0.2, (3) ≥ 1,000 UMIs, (4) ≥ 500 genes, and (5) ≤20% mitochondrial reads in each modality. Additionally, doublets were removed in both modalities using scDblFinder^52^, and ambient RNA was removed from the gene counts using SoupX^53^, yielding 108,220 cells. After integration across samples with Harmony^54^ on RNA-seq SCTransform^55^ principal components and on ATAC-seq latent semantic indexing (LSI) components, followed by cross-modality clustering using Seurat’s FindMultiModalNeighbors^56^, we annotated the clusters using canonical marker genes to obtain healthy proximal tubule (PT; 35,745 cells) and injured proximal tubule (PT-VCAM1; 8,730) cells. In all three datasets, peaks were called per cell type with MACS2^57^ via Signac’s CallPeaks^18^ using fragments pooled across all samples (standard chromosomes only, with ENCODE hg38 blacklist regions removed^58^; yielding 263,792 peaks in bone marrow, 380,154 in brain, and 368,013 in kidney).

### Features contributing to CREMAP latent factors

We evaluated the contribution of several per-cell measurements to the latent factors identified by CREMAP. Gene depth and peak depth were defined as each cell’s total gene and peak counts across the assay whereas genes detected and peaks detected were defined as the numbers of non-zero features in each modality (genes or peaks). S-phase and G2/M-phase scores were computed with Seurat’s CellCycleScoring^59^ using the cc.genes.updated.2019 gene list^60^, calculated as the mean expression of the cell-cycle phase-specific genes minus that of control genes matched by expression bins. For GM12878, TSS enrichment score, nucleosome signal, and the percentage of mitochondrial gene counts were also included as candidate covariates.

To evaluate the NF-κB transcriptional program in GM12878 (**Fig. S4b**), each cell’s activity for the Hallmark gene set of TNF-α signaling via NF-κB (gene set obtained from msigdbr^61^) was calculated using Seurat’s AddModuleScore on log-normalized gene counts with default parameters (24 expression bins, 100 control genes per set gene, random seed 1). The gene set was restricted to the 199 genes detected in the dataset, with *SERPINB9* excluded so that its own expression did not contribute to the score used to group cells. Cells were divided into tertiles at the 33rd and 67th percentiles of the score (low, 1,812 cells; mid, 1,811; high, 1,812). Because this score was derived from gene expression only, the resulting cell grouping was independent of chromatin accessibility.

### Peak–gene linking methods

For comparison, four existing peak–gene linking methods were evaluated alongside CREMAP: scMultiMap^24^ (v1.0.1), fasthurdle^17^ (v1.2.0; a hurdle model comprising a negative binomial count component and a binomial zero component), Signac^18^ (1.17.1) and FigR^19^ (0.1.0). Each method was run separately per cell type on all candidate pairs of a gene and a peak located within 250 kb of the gene’s transcription start site (TSS) yielding 754,339 candidate pairs in K562 and 881,135 in GM12878. In the multi-sample datasets, candidate pairs were further restricted to peaks accessible in the corresponding cell type. In the kidney dataset, genes and peaks were further limited to those detected in at least 1% of the cells within that cell type. CREMAP was run using 2 latent factors for consistency across datasets. In the single-sample datasets, no sample covariates were included, whereas in the multi-sample datasets, scMultiMap and fasthurdle were run both with and without the sample identity as a covariate. Specifically, sample adjustment was performed using the multi-sample option in scMultiMap, which estimates the mean of each feature per sample, and by including sample indicator variables in both the count and zero components of fasthurdle. Signac and FigR, which do not support sample covariates, were run without sample-level adjustment.

To call significant peak–gene links, we applied each method’s recommended multiple-testing correction procedure at a threshold of FDR < 0.05. For CREMAP and scMultiMap, the Benjamini– Hochberg procedure was applied across all tested candidate pairs. For Signac and FigR, two-sided P values were first recomputed from their reported one-sided P values and then adjusted using the Benjamini–Hochberg procedure. Finally, for fasthurdle, significant links were called using its two-stage Jiang–Doerge procedure^62^, while P values from the count and zero components were combined via Fisher’s method to generate a single ranking score for enrichment–recall analyses.

### Validation sets

Predicted peak–gene links were evaluated against two types of validation datasets: CRISPRi perturbation screens and eQTL datasets. For CRISPRi validation in K562, we used the training set of CRISPRi-tested element–gene pairs for ENCODE-rE2G^44^ (file crispr_data/EPCrisprBenchmark_combined_data.training_K562.GRCh38.tsv.gz, available at github.com/EngreitzLab/CRISPR_comparison). Each tested pair was labeled as “Regulated” if perturbation of the candidate element significantly affected target gene expression. Pairs labeled as “Regulated” were defined as positive pairs, whereas all remaining tested but non-significant pairs were defined as negative pairs. A candidate or predicted peak–gene link was matched to a tested element–gene pair if it targeted the same gene and at least half of the peak overlapped the tested element.

For eQTL validation, a candidate or predicted peak–gene link was matched to an eQTL pair if the SNP within the peak and the eGene matched the linked gene. Positive and negative SNP–gene pairs were defined using fine-mapping or multiple-testing results across three primary sources. First, from eQTL Catalogue release 7^11^, we downloaded SuSiE credible sets^63^ and full summary statistics from the eQTL Catalogue FTP server (ftp.ebi.ac.uk/pub/databases/spot/eQTL) across sorted cells, single cells, and bulk tissues (Geuvadis^64^; OneK1K^65^; BLUEPRINT^66^; GTEx^10^; ROSMAP^67^; CEDAR^68^; Schmiedel et al.^69^; Young et al.^70^). Within these datasets, a SNP–gene pair was defined as positive if the SNP was the lead SNP of a credible set with posterior inclusion probability (PIP) ≥ 0.2 or lay outside any credible set with P < 10^-6^, and as negative if it had PIP < 0.01 (or was absent from all credible sets) and P > 0.5. Second, for Brain eQTLs^71^, summary statistics for seven brain cell types from single-nucleus RNA sequencing of the dorsolateral prefrontal cortex in 424 ROSMAP participants were downloaded from Synapse (https://doi.org/10.7303/syn52335732; syn52335807). A pair was defined as positive if it was significant by the study’s two-step FDR, and as negative if it was not significant for the same gene with P > 0.5. Third, for kidney eQTLs, two cohorts fine-mapped with DAP-G^72,73^ were used. NEPTUNE tubulointerstitial (TUBE) and glomerular (GLOM) *cis*-eQTLs were fine-mapped following Han et al.^37^, using distance to the TSS as the prior. For GTEx v8 kidney cortex^10^, DAP-G fine-mapping results and nominal associations were downloaded from the GTEx Portal (gtexportal.org). A pair was defined as positive if its cluster PIP was > 0.8 and its SNP PIP > 0.5, and as negative if its cluster PIP was < 0.2 and its nominal P > 0.5. Only genes harboring both positive and negative pairs were included in the validation sets.

Each cell type was evaluated against one primary eQTL validation set matched to its biological identity. Specifically, GM12878 was compared against Geuvadis lymphoblastoid cell lines. Bone marrow cell types were benchmarked against BLUEPRINT monocytes (CD14+ monocytes), Fairfax^74^ (B cells), Kasela CD4 and CD8^75^ (CD4+ and CD8+ T cells) and Gilchrist^76^ (NK cells), all obtained from the eQTL Catalogue. Brain cell types were matched to the dataset of the corresponding cell type from Fujita et al. study. Lastly, kidney PT and PT-VCAM1 cells were compared against NEPTUNE TUBE, with GTEx v8 kidney cortex serving as a second kidney cohort (**Fig. S5a**). To evaluate robustness across additional datasets, each multi-sample cell type was further benchmarked against all remaining sets matching its cell type or tissue (38 additional sets, yielding 51 validation sets in total when combined with the 13 primary sets; **Fig. S8**). Similarly, GM12878 was evaluated against five additional sets from the eQTL Catalogue, comprising GTEx v8 EBV-transformed lymphocytes, GTEx v8 whole blood, and OneK1K naive, memory, and intermediate B cells (**Fig. 2b**).

### Validation metrics

The comparison analyses were performed separately for each cell type and validation set, restricted to genes tested by all evaluated peak–gene linking methods. We assessed the performance using two main metrics: area under the enrichment–recall curve (enrichment–recall AUC, which equals 1 for a random ranking) and recall at FDR < 0.05 (the fraction of positive validation pairs matched by at least one significant predicted link). Each validation pair was assigned the score of the best-scoring link from a given method, and all validation pairs were ranked by these scores. At each distinct score *c*, let TP(*c*) denote the number of positive pairs and *n*(*c*) denote the number of all validation pairs scoring at least as well as *c*, out of *P* positive and *N* validation pairs (both positive and negative pairs, including pairs not scored by the method). Recall and enrichment at threshold *c* were then defined as

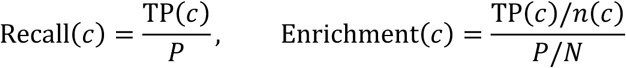

The enrichment–recall AUC was computed via trapezoidal rule as the area under the curve from recall 0 (anchored at the enrichment of the top-scoring pairs), to recall 1 and enrichment 1, with unscored validation pairs ordered randomly. To quantify uncertainty, 95% confidence intervals were calculated using 1,000 bootstrap resampling of genes for the enrichment–recall AUC and exact binomial intervals for recall. For stratified enrichment analyses, candidate pairs were partitioned into quartiles based on the gene’s and of the peak’s mean raw count per cell, and into five bins of peak–TSS distance (0–5, 5–20, 20–50, 50–100 and 100–250 kb). Then, the enrichment–recall AUC was computed within each stratum (**Figs. S2 and S7**). Finally, recall-at-k curves quantified the percentage of the eQTL-supported PT pairs (NEPTUNE TUBE) recovered among each method’s top *N* predicted pairs, ranked by P value (**Fig. S9a**; **Table S3**).

### *Trans*-pair test for inflation of correlation

In each cell type, 20,000 pairs of peaks and genes on different chromosomes were sampled from the 5,000 genes and 50,000 peaks with the highest mean counts, using gene expression and chromatin accessibility values log-normalized by each cell’s total counts. A pair was called correlated if the absolute value of its Pearson correlation across cells exceeded the 95^th^ percentile of an empirical null distribution generated by permuting cells between the gene and peak matrices (two-sided empirical P < 0.05; five permutations), so 5% of pairs are expected false positives.

To test whether this shared correlation was concentrated in a few leading components, the top 20 singular values of the matrix of peak–gene correlations were computed across the same genes and peaks using irlba 2.3.7. Each singular value was then normalized by its null expectation in the absence of shared variation, calculated as the mean singular value of the same rank across three permuted matrices in which the cell labels of the gene matrix were shuffled relative to the peak matrix (observed / expected; **Fig. S1c**).

To evaluate how effectively different adjustments mitigate this inflation, we repeated the trans-pair test after regressing out either CREMAP’s two latent factors (by removing each cell’s gene factor scores from the expression matrix and peak factor scores from the accessibility matrix; **Figs. 2a and 4f**) or sample identity (modeled as per-sample indicator variables that mimic a sample covariate; **Fig. 4f**). Specifically, letting *c*_none_ and *c*_a_ denote the fractions of trans pairs called correlated before and after adjustment *a*, respectively, the proportion of excess inflation removed was defined as (*c*_none_ − *c*_a_) / (*c*_none_ − 0.05), which equals 0 when the adjustment has no effect and 1 when inflation is reduced to the 5% null expectation.

### Characterization of the latent factors

To quantify which known sources of variation explain each latent factor (**Fig. 2d**; **Fig. S10a**), we separately regressed each factor’s gene and peak factor scores on the *q* per-cell measurements using the linear model 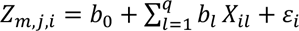 for factor *j*, modality *m* (gene or peak) and cell *i*. Letting *R*^2^(*S*) denote the proportion of variance explained by a model with a subset *S* of covariates, the variance contribution of covariate *l* was calculated as its Shapley value

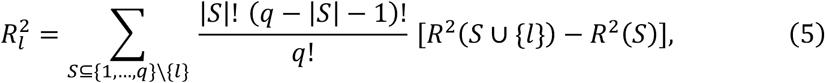

representing the average gain in *R*^2^ from adding covariate *l* across all possible entry orders of the covariates (Lindeman–Merenda–Gold decomposition^77^). These individual contributions sum to the full-model *R*^2^, with 1 − *R*^2^ reported as unexplained variance. Equation (5) was computed exactly, across all 2*^q^* covariate subsets. In kidney PT cells, sample identity and disease condition (CKD versus healthy) were included as additional covariates. Sample identity entered as a single grouped covariate (a set of sample indicator variables), and because disease condition is constant within each sample, the variance jointly explained by sample identity and disease condition was partitioned between them via equation (5).

To visualize factor activity across cells and summarize factor variation across donors (**Figs. S3a and S10a**), factor scores were projected onto weighted-nearest-neighbor (WNN) UMAP embeddings computed with Seurat’s FindMultiModalNeighbors and RunUMAP from RNA-seq principal components and ATAC-seq LSI components. Because latent factor signs are arbitrary, each factor was oriented to correlate positively with its strongest contributing covariate in cell lines or with gene sequencing depth in kidney PT cells. In kidney PT cells, each sample’s mean gene factor score was compared with its mean log(1 + gene depth) using Pearson correlation across the 23 samples, and differences between healthy and CKD samples were assessed using a two-sided Wilcoxon rank-sum test.

To identify biological pathways associated with each latent factor (**Figs. S3b, c and S10b**; **Table S2**), genes were ranked by their loadings on each factor, oriented consistently with the sign of the factor scores, and tested against the 50 MSigDB Hallmark gene sets using preranked gene set enrichment analysis^78^ (fgsea^79^; sets of 10 to 500 genes; 10,000 permutations). FDR values were computed separately within each factor. Gene sets were grouped according to the Hallmark process categories defined by Liberzon et al.^61^, and all gene sets within selected categories were shown.

### Genotypes and allele-specific chromatin accessibility

Genotypes were called from the ATAC-seq reads of each of the 23 kidney samples aligned to the GRCh38 reference genome using the SALSA pipeline^80^. Specifically, base-quality score recalibration and per-sample variant calling were performed with GATK^81^ (v4.2.0.0; HaplotypeCaller, GenotypeGVCFs, CNNScoreVariants, and FilterVariantTranches), while reads overlapping variants were remapped using WASP (v0.3.4)^82^ to eliminate reference mapping bias.

To identify variants exhibiting allele-specific chromatin accessibility (ASCA), allelic counts at biallelic heterozygous single-nucleotide variants (SNPs) were obtained using GATK ASEReadCounter from the PT cells of each sample without phasing. For each sample and each heterozygous SNP within a peak covered by ≥10 allelic reads with <10% of reads carrying other bases, allelic imbalance was evaluated using a two-sided binomial test against an expected proportion of 0.5 (518,589 tests across 62,297 peaks). Multiple-testing correction was performed using the Benjamini–Hochberg FDR across all tests, with significance defined at FDR < 0.05. Each tested SNP was classified into one of four tiers based on cross-sample consistency, namely ASCA in ≥ 3 samples (significant in ≥3 samples with a consistent allelic direction), ASCA in 1–2 samples (significant in one or two samples with a consistent direction), inconsistent (significant in opposite directions across samples and therefore excluded), or non-ASCA (tested but not significant in any sample). Each peak was assigned the highest tier among its respective SNPs, and a single lead SNP was selected per peak. At ASCA peaks, the lead SNP was chosen as the variant in the highest tier, breaking ties by the largest number of samples with significant allelic imbalance and then by the lowest FDR, whereas at non-ASCA peaks, the lead SNP was defined as the variant heterozygous in the largest number of samples (**Fig. 3e; Fig. S9b**).

To evaluate genotype effects across samples, peak and gene counts from PT cells were aggregated per sample (pseudobulk profiles). Sample H03 (156 PT cells and fewer than 1 million ATAC-seq reads) was excluded, and the remaining 22 samples were normalized using the trimmed mean of M-values (TMM) method^83^ and transformed to log_2_ counts per million (CPM) in edgeR^84^ with a prior count of 2. The genotype at each peak’s lead SNP was encoded as alternative allele dosage, defined as 1 or 2 based on the sample’s variant calls, 0 if no variant was called at a site covered by ≥8 reads, and missing otherwise. The genotype effect on a peak or gene was estimated as the ordinary least-squares slope of its log_2_ CPM against allele dosage, computed whenever ≥8 samples had non-missing genotypes and ≥2 genotype classes each contained ≥2 samples (**Fig. 3f**; **Fig. S5b**). For gene-level analyses, each peak was paired with its strongest lost link, and links were classified as positive or negative according to the sign of their association statistic (**Fig. S5c, d**).

### Downstream enrichment and concordance analyses

Unless stated otherwise, binary outcomes were modeled using logistic regression. Group estimates were computed on the log-odds scale at the mean of all covariates as estimated marginal means (emmeans) with Wald 95% confidence intervals, with each group compared against a designated reference group.

To evaluate eQTL enrichment across link categories (**Figs. 3d, 3g, and 4c**; **Figs. S5a, S6c, and S6f**), whether a labeled pair was eQTL-supported was modeled as a function of its link category, with non-linked pairs serving as the reference group, while adjusting for log(1 + distance to the nearest TSS), log(1 + peak depth), log(peak width) and log(1 + gene depth). When evaluating lost links specifically, the predictor category was defined as the ASCA tier of the peak, with non-ASCA peaks serving as the reference group.

To assess link calling at ASCA peaks (**Figs. 3e and 4d**; **Figs. S6d and S9b**), each tested pair overlapping a peak with a testable SNP was included once per method, and significance status was modeled as a function of the peak’s ASCA tier, the linking method (or run), and their interaction. Models were adjusted for log(1 + allelic reads in the peak), log(1 + number of allelic tests), log(1 + number of heterozygous samples), log(1 + peak depth), log(peak width) and log(1 + gene depth). For each ASCA tier, the change in log odds between unadjusted and adjusted runs of a given method was tested against the corresponding change at non-ASCA peaks. Similarly, to compare across methods, the log odds ratio of each ASCA tier relative to non-ASCA peaks was estimated for each method, and each competing method was compared against CREMAP via the tier × method interaction term.

To test directional concordance between ASCA and cross-sample genotype effects (Fig. S5b, c), whether the allelic direction of ASCA and the pseudobulk genotype effect shared the same sign was modeled as a function of the peak’s ASCA tier, with one observation per peak. Models were adjusted for log(1 + allelic reads at the lead SNP), log(1 + number of heterozygous samples), log(1 + number of allelic tests), log(1 + peak depth) and log(peak width), as well as log(1 + gene depth) when modeling genotype effects on gene expression.

Finally, to compare how well CREMAP retains validated versus non-validated links identified by competing methods, the proportions of links also called by CREMAP were compared using two-sided Fisher’s exact tests (**Fig. S2e, f**).

### Estimation of genotype effect absorbed by latent factors

Because all cells from a given sample share the same genotype, a latent factor can capture a genetic effect only through its between-sample variation. To measure how much of a SNP’s genotype variation across samples is captured by the latent factors, each factor’s gene and peak scores were averaged across the cells within each sample, and the alternative allele genotype dosage *G_vs_* of lead SNP *v* in sample *s* was regressed on these per-sample mean scores *η̂_s_*_j_ for the first *k* factors (*k* = 1 to 20, evaluated separately for gene and peak factors), with each sample *s* weighted by its cell count,

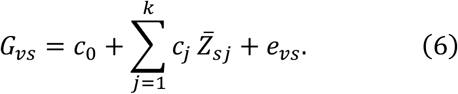

The weighted coefficient of determination *R*^2^(*k*) from equation (6) represents the proportion of the genotype variance for SNP *v* explained by the first k factors. It was computed across the 23,317 lead SNPs genotyped in all 23 kidney PT samples and summarized by its median and quartiles across SNPs, generating curves that describe the proportion of genotype variance explained as a function of numbers of latent factors (**Fig. 4g**). The null expectation by chance was estimated in the same way after randomly permuting genotype dosages across samples (20 permutations), whereas including sample identity as a covariate accounts for 100% of the genotype variance by construction.

### Software

Analyses used R 4.3.3 with Seurat 5.2.1, SeuratObject 5.0.2, Signac 1.17.1, harmony 1.2.0, FigR 0.1.0, fasthurdle 1.2.0, scMultiMap 1.0.1, CREMAP 0.1.0, fgsea 1.28.0 with the MSigDB Hallmark sets of msigdbr 26.1.0, edgeR 4.0.16, irlba 2.3.7, sandwich 3.1-1, emmeans 2.0.0, GenomicRanges 1.54.1, ensembldb 2.26.0 with EnsDb.Hsapiens.v86, data.table 1.18.4, ggplot2 4.0.3, ggforce 0.4.2, ggtext 0.1.2 and patchwork 1.3.2. Bone-marrow fragment files were generated with sinto 0.10.1, peaks were called with MACS2 2.2.4, and genotyping used GATK 4.2.0.0, BWA 0.7.17, WASP 0.3.4, bcftools 1.24 and samtools 1.24. Kidney quality control, doublet removal and integration used SoupX 1.6.2, scDblFinder 1.16.0, SnapATAC2 2.8.0, scanpy 1.10.3, Seurat 5.3.0 with sctransform 0.4.2 and glmGamPoi 1.14.0, Signac 1.14.0 and harmony 1.2.3.

### Data availability

K562: ArrayExpress E-MTAB-11264 (sample ERS12227353). GM12878: GEO GSE166797. Bone marrow: GEO GSE194122 (fragments regenerated from ENA SRR17693253 to SRR17693265). Brain: GEO GSE214979. Kidney: GEO GSE232222^50^ and GEO GSE220289^51^. The five pediatric kidney samples generated in our laboratory are part of a study in preparation; they will be deposited in GEO with that study and are available from the corresponding author upon reasonable request.

### Code availability

CREMAP is available as an R package at https://github.com/dongwonlee-lab/CREMAP. Code for the analyses is available upon request.

## Acknowledgments

We acknowledge the Boston Children’s Hospital High-Performance Computing (BCH HPC) resources, including the E3 cluster, for supporting the computational analyses performed in this study. We also thank the members of the Sampson and Lee labs for their valuable discussions, feedback, and support throughout the course of this work. During manuscript preparation, the authors used large language models (LLMs) to improve the clarity and phrasing of the text. All suggestions were reviewed and edited by the authors, who take full responsibility for the content of the manuscript.

## Author contributions

Conceptualization: DL, NDN

Data curation: GK, MTM, NDN, SKH

Formal analysis: NDN

Methodology: DL, NDN

Investigation: DL, NDN

Visualization: NDN

Funding acquisition: DL, MGS

Project administration: DL, NDN

Resources: MGS, NDN

Software: NDN

Validation: NDN

Supervision: DL

Writing – original draft: DL, NDN

Writing – review & editing: DL, GK, MGS, MTM, NDN, SKH

## Ethics statement

This study analyzed publicly available, de-identified single-cell multiome datasets. In addition, the five newly generated kidney samples were analyzed under a protocol reviewed by the Institutional Review Board of Boston Children’s Hospital (IRB-P00033405).

## References

1. Visscher, P. M. et al. 10 years of GWAS discovery: biology, function, and translation. Am. J. Hum. Genet. 101, 5–22 (2017). 10.1016/j.ajhg.2017.06.005

2. Sollis, E. et al. The NHGRI-EBI GWAS Catalog: knowledgebase and deposition resource. Nucleic Acids Res. 51, D977–D985 (2023). 10.1093/nar/gkac1010

3. Maurano, M. T. et al. Systematic localization of common disease-associated variation in regulatory DNA. Science 337, 1190–1195 (2012). 10.1126/science.1222794

4. Mostafavi, H., Spence, J. P., Naqvi, S. & Pritchard, J. K. Systematic differences in discovery of genetic effects on gene expression and complex traits. Nat. Genet. 55, 1866–1875 (2023). 10.1038/s41588-023-01529-1

5. Claussnitzer, M. et al. A brief history of human disease genetics. Nature 577, 179–189 (2020). 10.1038/s41586-019-1879-7

6. Gamazon, E. R. et al. Using an atlas of gene regulation across 44 human tissues to inform complex disease- and trait-associated variation. Nat. Genet. 50, 956–967 (2018). 10.1038/s41588-018-0154-4

7. Gazal, S. et al. Combining SNP-to-gene linking strategies to identify disease genes and assess disease omnigenicity. Nat. Genet. 54, 827–836 (2022). 10.1038/s41588-022-01087-y

8. Strober, B. J. et al. Dynamic genetic regulation of gene expression during cellular differentiation. Science 364, 1287–1290 (2019). 10.1126/science.aaw0040

9. Nathan, A. et al. Single-cell eQTL models reveal dynamic T cell state dependence of disease loci. Nature 606, 120–128 (2022). 10.1038/s41586-022-04713-1

10. GTEx Consortium. The GTEx Consortium atlas of genetic regulatory effects across human tissues. Science 369, 1318–1330 (2020). 10.1126/science.aaz1776

11. Kerimov, N. et al. A compendium of uniformly processed human gene expression and splicing quantitative trait loci. Nat. Genet. 53, 1290–1299 (2021). 10.1038/s41588-021-00924-w

12. Heinz, S., Romanoski, C. E., Benner, C. & Glass, C. K. The selection and function of cell type-specific enhancers. Nat. Rev. Mol. Cell Biol. 16, 144–154 (2015). 10.1038/nrm3949

13. Chen, S., Lake, B. B. & Zhang, K. High-throughput sequencing of the transcriptome and chromatin accessibility in the same cell. Nat. Biotechnol. 37, 1452–1457 (2019). 10.1038/s41587-019-0290-0

14. Ma, S. et al. Chromatin potential identified by shared single-cell profiling of RNA and chromatin. Cell 183, 1103–1116 (2020). 10.1016/j.cell.2020.09.056

15. Vandereyken, K., Sifrim, A., Thienpont, B. & Voet, T. Methods and applications for single-cell and spatial multi-omics. Nat. Rev. Genet. 24, 494–515 (2023). 10.1038/s41576-023-00580-2

16. Zhang, K. et al. A single-cell atlas of chromatin accessibility in the human genome. Cell 184, 5985–6001 (2021). 10.1016/j.cell.2021.10.024

17. Kanai, M. et al. Population-scale immune multiome atlas reveals regulatory disease mechanisms. Nature (2026). 10.1038/s41586-026-11078-2

18. Stuart, T., Srivastava, A., Madad, S., Lareau, C. A. & Satija, R. Single-cell chromatin state analysis with Signac. Nat. Methods 18, 1333–1341 (2021). 10.1038/s41592-021-01282-5

19. Kartha, V. K. et al. Functional inference of gene regulation using single-cell multi-omics. Cell Genom. 2, 100166 (2022). 10.1016/j.xgen.2022.100166

20. Granja, J. M. et al. ArchR is a scalable software package for integrative single-cell chromatin accessibility analysis. Nat. Genet. 53, 403–411 (2021). 10.1038/s41588-021-00790-6

21. Fang, R. et al. Comprehensive analysis of single cell ATAC-seq data with SnapATAC. Nat. Commun. 12, 1337 (2021). 10.1038/s41467-021-21583-9

22. Sakaue, S. et al. Tissue-specific enhancer-gene maps from multimodal single-cell data identify causal disease alleles. Nat. Genet. 56, 615–626 (2024). 10.1038/s41588-024-01682-1

23. Liu, H. et al. Kidney multiome-based genetic scorecard reveals convergent coding and regulatory variants. Science 387, eadp4753 (2025). 10.1126/science.adp4753

24. Su, C., Lee, D., Jin, P. & Zhang, J. scMultiMap: cell-type-specific mapping of enhancers and target genes from single-cell multimodal data. Nat. Commun. 16, 3941 (2025). 10.1038/s41467-025-59306-z

25. Dorans, E., Jagadeesh, K., Dey, K. & Price, A. L. Linking regulatory variants to target genes by integrating single-cell multiome methods and genomic distance. Nat. Genet. 57, 1649–1658 (2025). 10.1038/s41588-025-02220-3

26. Pliner, H. A. et al. Cicero predicts cis-regulatory DNA interactions from single-cell chromatin accessibility data. Mol. Cell 71, 858–871 (2018). 10.1016/j.molcel.2018.06.044

27. Sheth, M. U. et al. Mapping enhancer-gene regulatory interactions from single-cell data. Nat. Genet. 58, 1941–1952 (2026). 10.1038/s41588-026-02695-8

28. Fulco, C. P. et al. Activity-by-contact model of enhancer-promoter regulation from thousands of CRISPR perturbations. Nat. Genet. 51, 1664–1669 (2019). 10.1038/s41588-019-0538-0

29. Buettner, F. et al. Computational analysis of cell-to-cell heterogeneity in single-cell RNA-sequencing data reveals hidden subpopulations of cells. Nat. Biotechnol. 33, 155–160 (2015). 10.1038/nbt.3102

30. Ilicic, T. et al. Classification of low quality cells from single-cell RNA-seq data. Genome Biol. 17, 29 (2016). 10.1186/s13059-016-0888-1

31. Tung, P.-Y. et al. Batch effects and the effective design of single-cell gene expression studies. Sci. Rep. 7, 39921 (2017). 10.1038/srep39921

32. Zimmerman, K. D., Espeland, M. A. & Langefeld, C. D. A practical solution to pseudoreplication bias in single-cell studies. Nat. Commun. 12, 738 (2021). 10.1038/s41467-021-21038-1

33. Degner, J. F. et al. DNase I sensitivity QTLs are a major determinant of human expression variation. Nature 482, 390–394 (2012). 10.1038/nature10808

34. Leek, J. T. & Storey, J. D. Capturing heterogeneity in gene expression studies by surrogate variable analysis. PLoS Genet. 3, 1724–1735 (2007). 10.1371/journal.pgen.0030161

35. Stegle, O., Parts, L., Piipari, M., Winn, J. & Durbin, R. Using probabilistic estimation of expression residuals (PEER) to obtain increased power and interpretability of gene expression analyses. Nat. Protoc. 7, 500–507 (2012). 10.1038/nprot.2011.457

36. Parsana, P. et al. Addressing confounding artifacts in reconstruction of gene co-expression networks. Genome Biol. 20, 94 (2019). 10.1186/s13059-019-1700-9

37. Han, S. K. et al. Mapping genomic regulation of kidney disease and traits through high-resolution and interpretable eQTLs. Nat. Commun. 14, 2229 (2023). 10.1038/s41467-023-37691-7

38. Schulz, W. W., Hagler, H. K., Buja, L. M. & Erdös, E. G. Ultrastructural localization of angiotensin I-converting enzyme (EC 3.4.15.1) and neutral metalloendopeptidase (EC 3.4.24.11) in the proximal tubule of the human kidney. Lab. Invest. 59, 789–797 (1988). PMID: 2848979

39. Rigat, B. et al. An insertion/deletion polymorphism in the angiotensin I-converting enzyme gene accounting for half the variance of serum enzyme levels. J. Clin. Invest. 86, 1343–1346 (1990). 10.1172/JCI114844

40. Lewis, E. J., Hunsicker, L. G., Bain, R. P. & Rohde, R. D. The effect of angiotensin-converting-enzyme inhibition on diabetic nephropathy. N. Engl. J. Med. 329, 1456–1462 (1993). 10.1056/NEJM199311113292004

41. The GISEN Group (Gruppo Italiano di Studi Epidemiologici in Nefrologia). Randomised placebo-controlled trial of effect of ramipril on decline in glomerular filtration rate and risk of terminal renal failure in proteinuric, non-diabetic nephropathy. Lancet 349, 1857–1863 (1997). 10.1016/S0140-6736(96)11445-8

42. Kang, H. M. et al. Defective fatty acid oxidation in renal tubular epithelial cells has a key role in kidney fibrosis development. Nat. Med. 21, 37–46 (2015). 10.1038/nm.3762

43. Lake, B. B. et al. An atlas of healthy and injured cell states and niches in the human kidney. Nature 619, 585–594 (2023). 10.1038/s41586-023-05769-3

44. Gschwind, A. R. et al. An encyclopedia of human enhancer-gene regulatory interactions. Nature 657, 179–190 (2026). 10.1038/s41586-026-10781-4

45. Tucker, L. R. An inter-battery method of factor analysis. Psychometrika 23, 111–136 (1958). 10.1007/BF02289009

46. Xu, W. et al. ISSAAC-seq enables sensitive and flexible multimodal profiling of chromatin accessibility and gene expression in single cells. Nat. Methods 19, 1243–1249 (2022). 10.1038/s41592-022-01601-4

47. Wu, K. E., Yost, K. E., Chang, H. Y. & Zou, J. BABEL enables cross-modality translation between multiomic profiles at single-cell resolution. Proc. Natl Acad. Sci. USA 118, e2023070118 (2021). 10.1073/pnas.2023070118

48. Luecken, M. D. et al. A sandbox for prediction and integration of DNA, RNA, and proteins in single cells. In Proc. Neural Information Processing Systems Track on Datasets and Benchmarks 1 (NeurIPS, 2021). https://datasets-benchmarks-proceedings.neurips.cc/paper/2021/hash/158f3069a435b314a80bdcb024f8e422-Abstract-round2.html

49. Anderson, A. G. et al. Single nucleus multiomics identifies ZEB1 and MAFB as candidate regulators of Alzheimer’s disease-specific cis-regulatory elements. Cell Genom. 3, 100263 (2023). 10.1016/j.xgen.2023.100263

50. Wilson, P. C. et al. Mosaic loss of Y chromosome is associated with aging and epithelial injury in chronic kidney disease. Genome Biol. 25, 36 (2024). 10.1186/s13059-024-03173-2

51. Ledru, N. et al. Predicting proximal tubule failed repair drivers through regularized regression analysis of single cell multiomic sequencing. Nat. Commun. 15, 1291 (2024). 10.1038/s41467-024-45706-0

52. Germain, P.-L., Lun, A., Garcia Meixide, C., Macnair, W. & Robinson, M. D. Doublet identification in single-cell sequencing data using scDblFinder. F1000Research 10, 979 (2021). 10.12688/f1000research.73600.2

53. Young, M. D. & Behjati, S. SoupX removes ambient RNA contamination from droplet-based single-cell RNA sequencing data. GigaScience 9, giaa151 (2020). 10.1093/gigascience/giaa151

54. Korsunsky, I. et al. Fast, sensitive and accurate integration of single-cell data with Harmony. Nat. Methods 16, 1289–1296 (2019). 10.1038/s41592-019-0619-0

55. Hafemeister, C. & Satija, R. Normalization and variance stabilization of single-cell RNA-seq data using regularized negative binomial regression. Genome Biol. 20, 296 (2019). 10.1186/s13059-019-1874-1

56. Hao, Y. et al. Integrated analysis of multimodal single-cell data. Cell 184, 3573–3587 (2021). 10.1016/j.cell.2021.04.048

57. Zhang, Y. et al. Model-based analysis of ChIP-Seq (MACS). Genome Biol. 9, R137 (2008). 10.1186/gb-2008-9-9-r137

58. Amemiya, H. M., Kundaje, A. & Boyle, A. P. The ENCODE Blacklist: identification of problematic regions of the genome. Sci. Rep. 9, 9354 (2019). 10.1038/s41598-019-45839-z

59. Hao, Y. et al. Dictionary learning for integrative, multimodal and scalable single-cell analysis. Nat. Biotechnol. 42, 293–304 (2024). 10.1038/s41587-023-01767-y

60. Tirosh, I. et al. Dissecting the multicellular ecosystem of metastatic melanoma by single-cell RNA-seq. Science 352, 189–196 (2016). 10.1126/science.aad0501

61. Liberzon, A. et al. The Molecular Signatures Database (MSigDB) hallmark gene set collection. Cell Syst. 1, 417–425 (2015). 10.1016/j.cels.2015.12.004

62. Jiang, H. & Doerge, R. W. A two-step multiple comparison procedure for a large number of tests and multiple treatments. Stat. Appl. Genet. Mol. Biol. 5, 28 (2006). 10.2202/1544-6115.1223

63. Wang, G., Sarkar, A., Carbonetto, P. & Stephens, M. A simple new approach to variable selection in regression, with application to genetic fine mapping. J. R. Stat. Soc. B 82, 1273–1300 (2020). 10.1111/rssb.12388

64. Lappalainen, T. et al. Transcriptome and genome sequencing uncovers functional variation in humans. Nature 501, 506–511 (2013). 10.1038/nature12531

65. Yazar, S. et al. Single-cell eQTL mapping identifies cell type-specific genetic control of autoimmune disease. Science 376, eabf3041 (2022). 10.1126/science.abf3041

66. Chen, L. et al. Genetic drivers of epigenetic and transcriptional variation in human immune cells. Cell 167, 1398–1414 (2016). 10.1016/j.cell.2016.10.026

67. Ng, B. et al. An xQTL map integrates the genetic architecture of the human brain’s transcriptome and epigenome. Nat. Neurosci. 20, 1418–1426 (2017). 10.1038/nn.4632

68. Momozawa, Y. et al. IBD risk loci are enriched in multigenic regulatory modules encompassing putative causative genes. Nat. Commun. 9, 2427 (2018). 10.1038/s41467-018-04365-8

69. Schmiedel, B. J. et al. Impact of genetic polymorphisms on human immune cell gene expression. Cell 175, 1701–1715 (2018). 10.1016/j.cell.2018.10.022

70. Young, A. M. H. et al. A map of transcriptional heterogeneity and regulatory variation in human microglia. Nat. Genet. 53, 861–868 (2021). 10.1038/s41588-021-00875-2

71. Fujita, M. et al. Cell subtype-specific effects of genetic variation in the Alzheimer’s disease brain. Nat. Genet. 56, 605–614 (2024). 10.1038/s41588-024-01685-y

72. Wen, X., Lee, Y., Luca, F. & Pique-Regi, R. Efficient integrative multi-SNP association analysis via deterministic approximation of posteriors. Am. J. Hum. Genet. 98, 1114– 1129 (2016). 10.1016/j.ajhg.2016.03.029

73. Lee, Y., Luca, F., Pique-Regi, R. & Wen, X. Bayesian multi-SNP genetic association analysis: control of FDR and use of summary statistics. Preprint at bioRxiv (2018). 10.1101/316471

74. Fairfax, B. P. et al. Genetics of gene expression in primary immune cells identifies cell type-specific master regulators and roles of HLA alleles. Nat. Genet. 44, 502–510 (2012). 10.1038/ng.2205

75. Kasela, S. et al. Pathogenic implications for autoimmune mechanisms derived by comparative eQTL analysis of CD4+ versus CD8+ T cells. PLoS Genet. 13, e1006643 (2017). 10.1371/journal.pgen.1006643

76. Gilchrist, J. J. et al. Natural killer cells demonstrate distinct eQTL and transcriptome-wide disease associations, highlighting their role in autoimmunity. Nat. Commun. 13, 4073 (2022). 10.1038/s41467-022-31626-4

77. Grömping, U. Relative importance for linear regression in R: the package relaimpo. J. Stat. Softw. 17, 1–27 (2006). 10.18637/jss.v017.i01

78. Subramanian, A., et al. Gene set enrichment analysis: a knowledge-based approach for interpreting genome-wide expression profiles. Proc. Natl Acad. Sci. USA 102, 15545–15550 (2005). 10.1073/pnas.0506580102

79. Korotkevich, G. et al. Fast gene set enrichment analysis. Preprint at bioRxiv (2021). 10.1101/060012

80. Wilson, P. C. et al. Multimodal single cell sequencing implicates chromatin accessibility and genetic background in diabetic kidney disease progression. Nat. Commun. 13, 5253 (2022). 10.1038/s41467-022-32972-z

81. McKenna, A. et al. The Genome Analysis Toolkit: a MapReduce framework for analyzing next-generation DNA sequencing data. Genome Res. 20, 1297–1303 (2010). 10.1101/gr.107524.110

82. van de Geijn, B., McVicker, G., Gilad, Y. & Pritchard, J. K. WASP: allele-specific software for robust molecular quantitative trait locus discovery. Nat. Methods 12, 1061– 1063 (2015). 10.1038/nmeth.3582

83. Robinson, M. D. & Oshlack, A. A scaling normalization method for differential expression analysis of RNA-seq data. Genome Biol. 11, R25 (2010). 10.1186/gb-2010-11-3-r25

84. Robinson, M. D., McCarthy, D. J. & Smyth, G. K. edgeR: a Bioconductor package for differential expression analysis of digital gene expression data. Bioinformatics 26, 139– 140 (2010). 10.1093/bioinformatics/btp616

