## Supplementary Document for "A method to disentangle enhancer-gene interactions from latent factors in single-cell multi-omics data"

### Supplementary figures and table legends

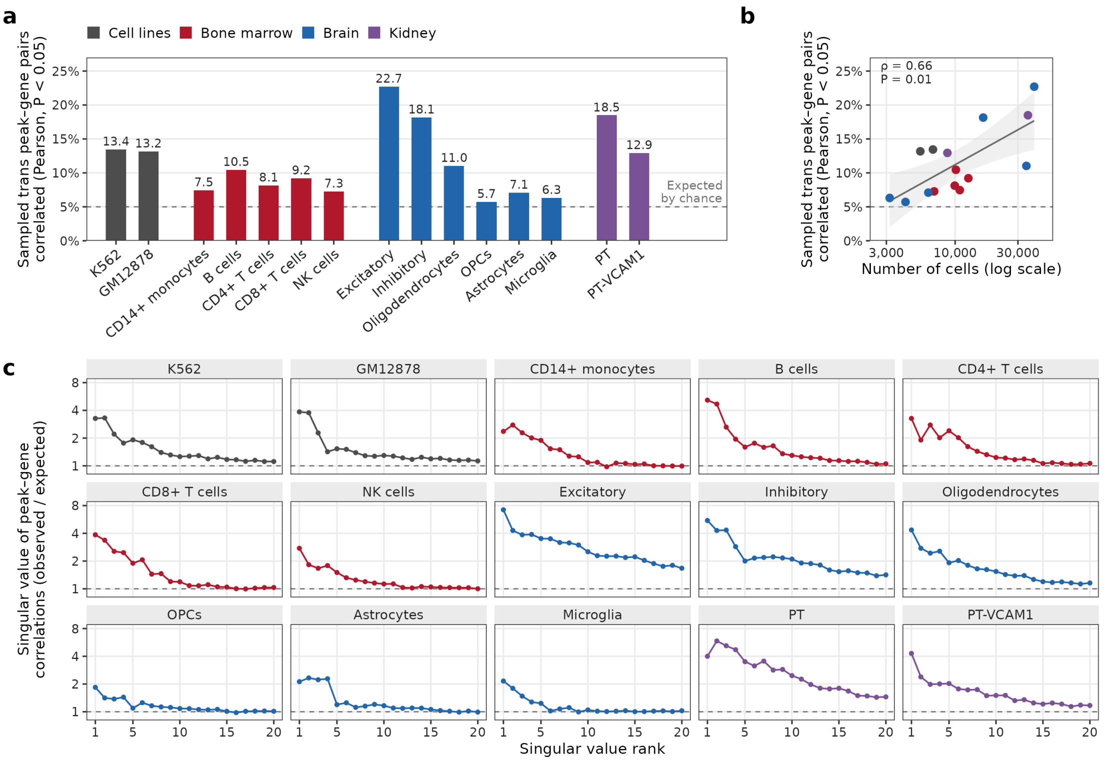


**Figure S1. Shared variation creates spurious peak–gene correlation in single-cell multiome data. (a)** Percentage of 20,000 randomly sampled *trans* pairs with significant Pearson correlation ($P < 0.05$), relative to the 5% null expectation (dashed line). **(b)** Percentages of significantly correlated *trans* pairs against cell counts across the 15 cell types, showing the least-squares fit with 95% confidence band and Spearman correlation (ρ). **(c)** Top 20 singular values of the peak–gene correlation matrix in the same cells (top 5,000 genes and 50,000 peaks by mean count), divided at each rank by the value expected without shared variation. The dashed line indicates 1, the null expectation. In (a)–(c), color indicates dataset.

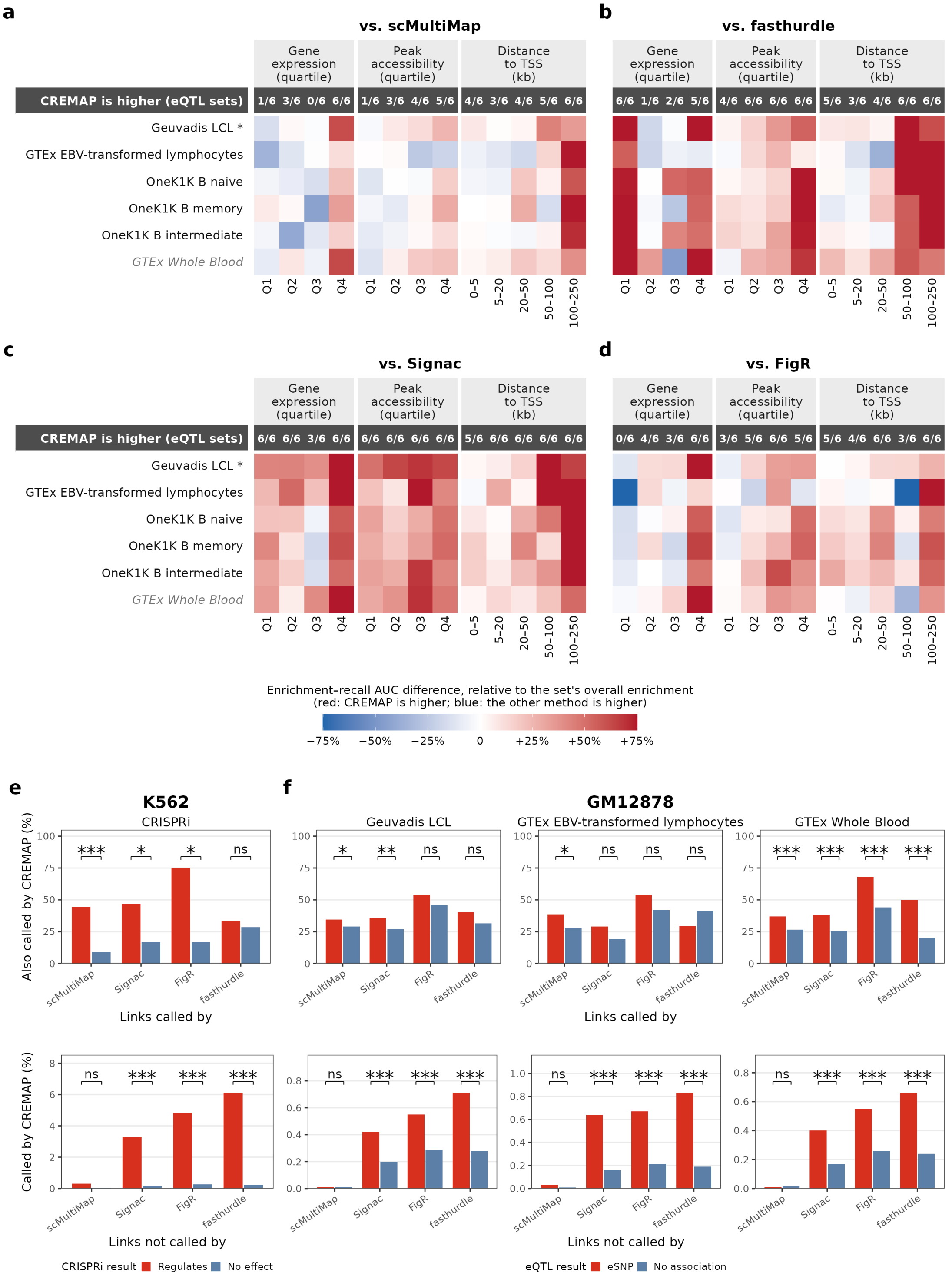


**Figure S2. Performance of CREMAP compared with other methods on single-sample cell line datasets across different contexts. (a)–(d)** Differences in enrichment–recall AUC between CREMAP and each competing method across candidate-pair strata (columns) for the six GM12878 eQTL sets (rows, with * marking the primary set), scaled to each set’s overall enrichment. The K562 CRISPRi set has too few positives per stratum and is excluded. **(e)** Comparison of significant links (FDR < 0.05) on K562 CRISPRi-tested pairs, stratified by CRISPRi result. Top: percentage of each method's called links also called by CREMAP. Bottom: percentage of tested pairs missed by each method that are called by CREMAP. **(f)** Same comparison for GM12878 across three eQTL sets, stratified by whether the peak contains a fine-mapped eSNP for the gene. In (e) and (f), analyses include pairs tested by both methods, and brackets denote two-sided Fisher's exact test significance (* P < 0.05; ** P < 0.01; *** P < 0.001; ns, not significant).

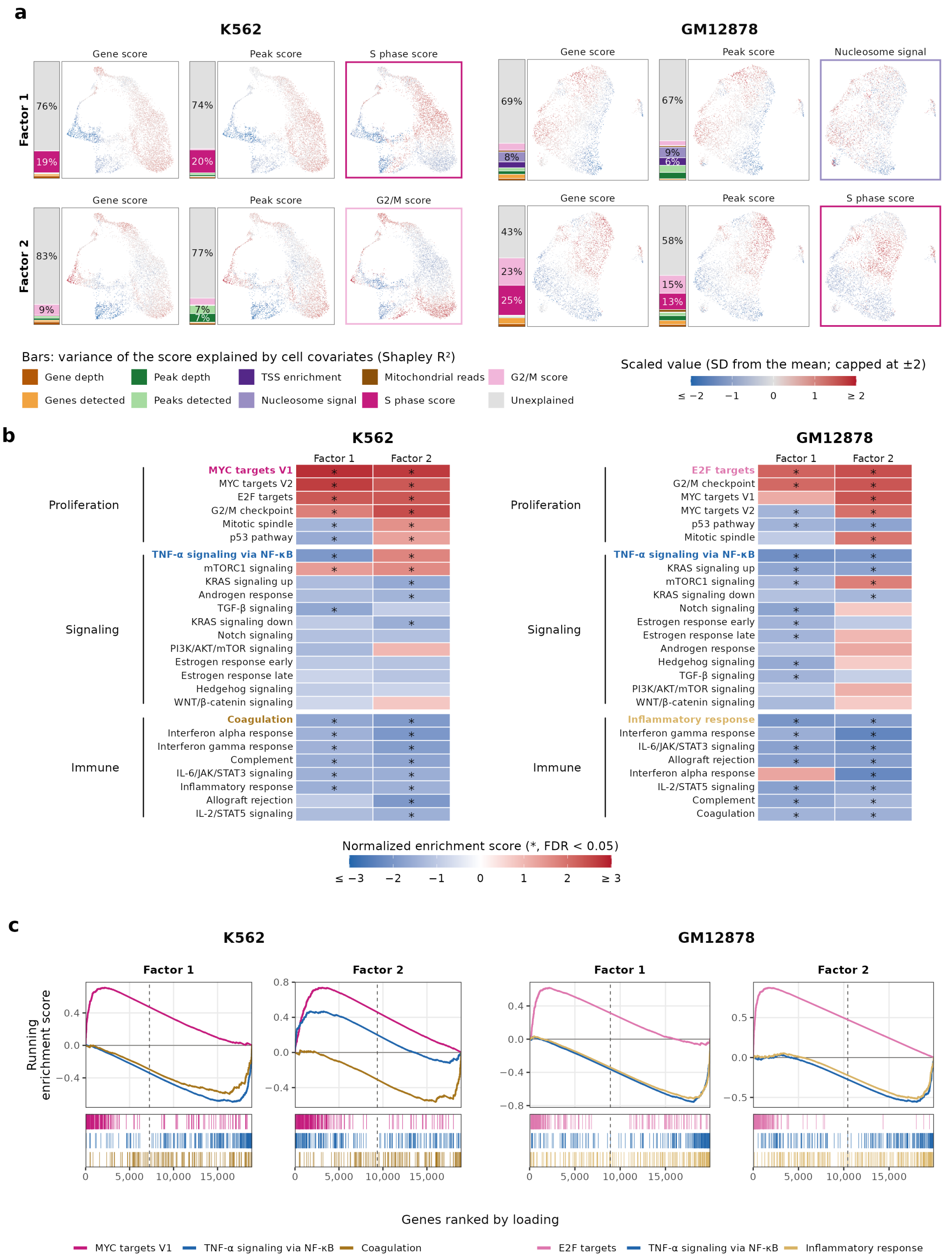


**Figure S3. Technical factors and biological signals captured by latent factors in single-sample cell line datasets.** K562 (left) and GM12878 (right) were analyzed in separate runs. **(a)** Each factor's gene score, peak score, and strongest covariate (outlined in its legend color) on the cells' weighted-nearest-neighbor UMAP. **(b)** Normalized enrichment score (NES) of Hallmark gene sets on each factor's gene loadings. **(c)** Running enrichment score of one representative set per category along genes ranked by loading. Ticks indicate genes in each set, and the dashed line indicates where loadings cross zero.

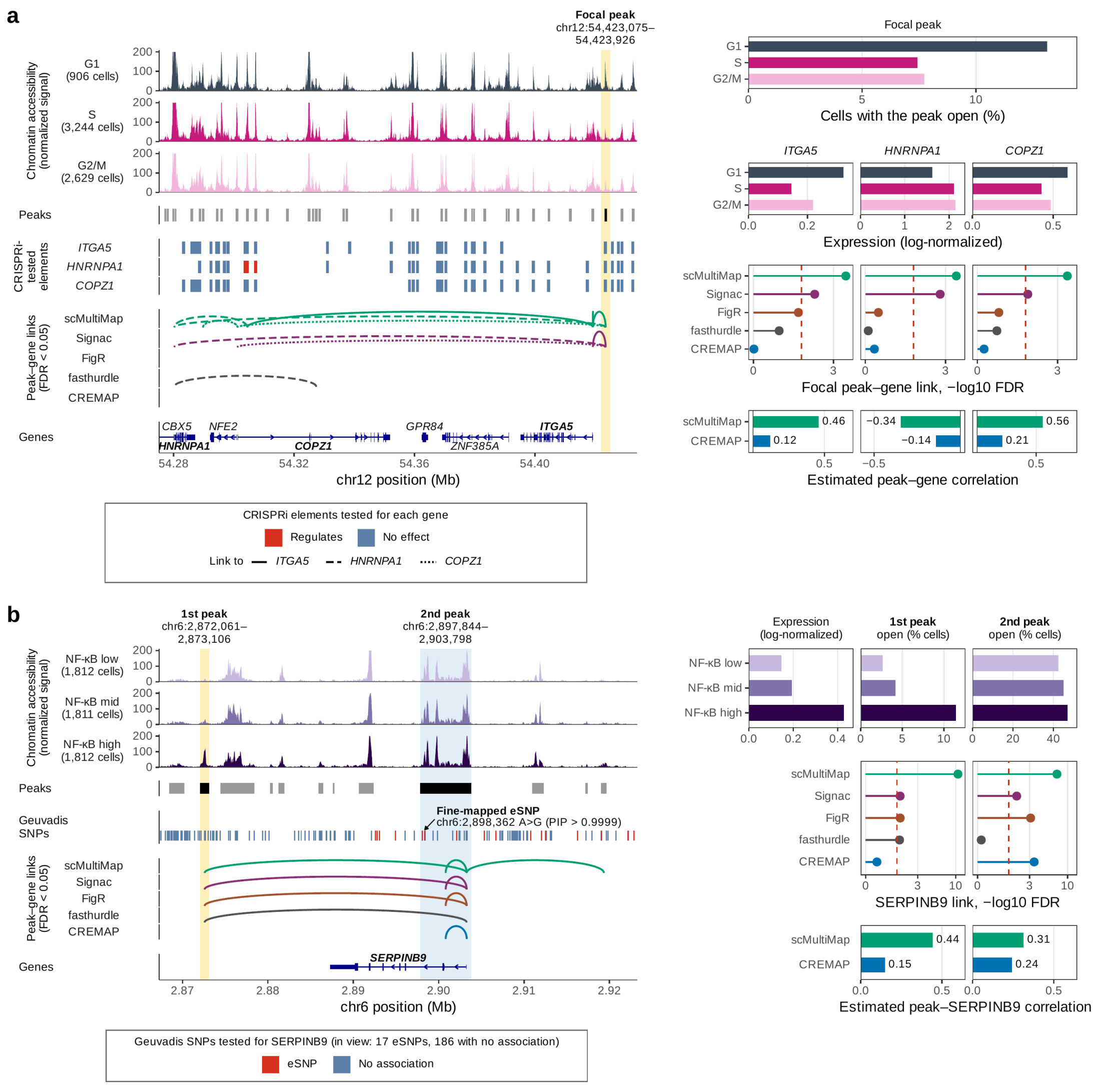


**Figure S4. CREMAP eliminates false-positive peak–gene links driven by shared variation.** **(a)** A focal peak (yellow) in K562 that scMultiMap links to *ITGA5*, *HNRNPA1*, and *COPZ1* (Signac also links this peak), none of which is confirmed by CRISPRi, whereas CREMAP removes all three links. **(b)** *SERPINB9* locus in GM12878, with cells grouped into tertiles of a TNF-α/NF-κB program score. The 1st peak (yellow) lacks eQTL support and is falsely linked by the four other methods, whereas the 2nd peak (blue) overlaps *SERPINB9*'s fine-mapped eSNP and promoter and is retained by CREMAP. In (a) and (b), left panels show chromatin accessibility, peaks, tested elements or SNPs, significant peak–gene links (FDR < 0.05), and genes, and right panels show peak accessibility and gene expression across cell states, link significance (red dashed line, FDR = 0.05), and estimated correlations.

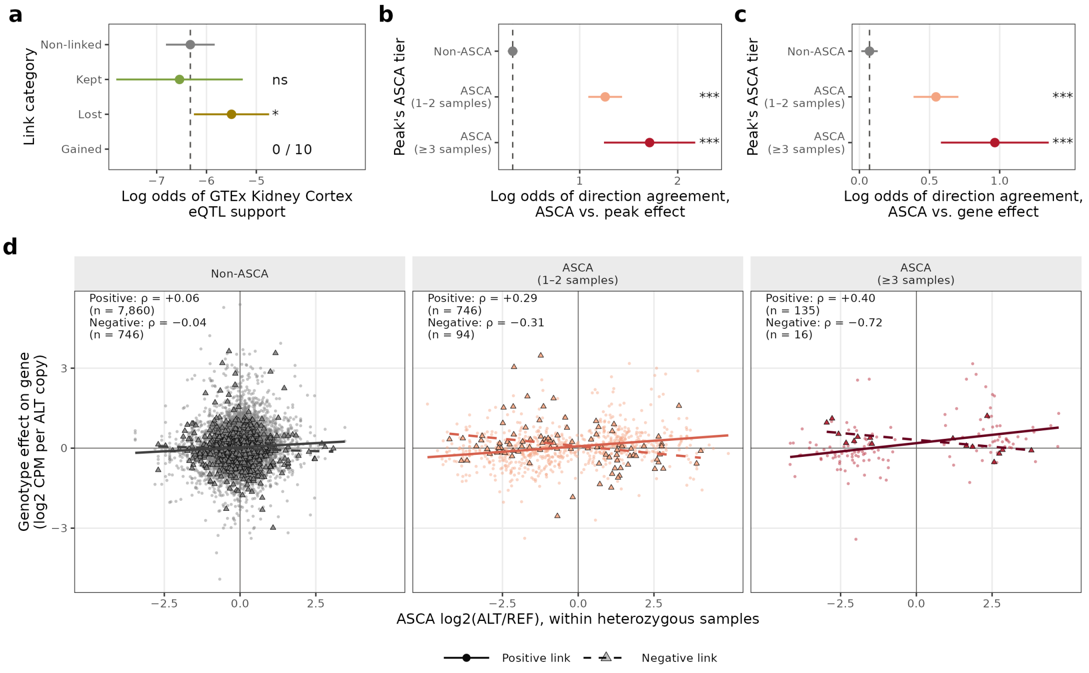


**Figure S5. Genotype effects of ASCA variants on the peak and the linked gene. (a)** Log odds of GTEx Kidney Cortex eQTL support by link category, with the dashed line showing the Non-linked reference. Categories with <3 eQTL-supported or unsupported links are given as a count (supported / tested). **(b)** For peaks that lost a link, log odds that ASCA and the genotype effect on the peak agree in direction, by the peak's ASCA tier (adjusted for allelic coverage and depth), with dashed line as Non-ASCA reference. **(c)** Log odds of direction agreement between ASCA and the genotype effect on linked gene expression (positive links), by the peak's ASCA tier. **(d)** For the strongest lost link of each peak, ASCA log2(ALT/REF) against the genotype effect on linked gene expression, by the peak's ASCA tier. Lines show least-squares fits for positive and negative links, and ρ indicates Spearman correlation across *n* links. In (a)–(c), dots and whiskers show estimates with 95% CI, and asterisks denote significance versus the reference (* $P < 0.05$; ** $P < 0.01$; *** $P < 0.001$; ns, not significant).

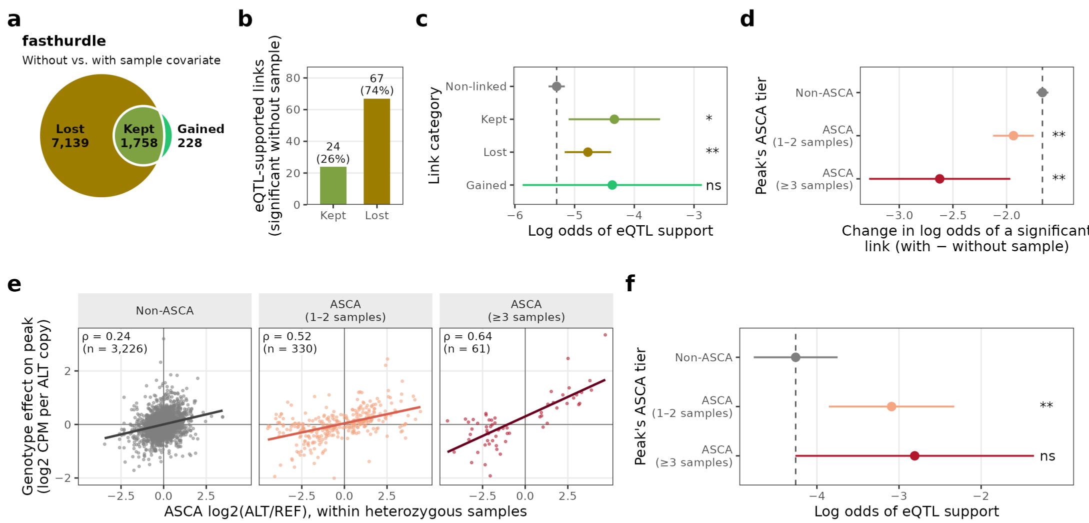


**Figure S6. Including a sample covariate in a regression-based model removes genotype-driven and eQTL-supported links.** Panels (a)–(f) show fasthurdle results in kidney proximal tubule (PT) cells, with significant links called using the two-stage Jiang–Doerge rule (FDR < 0.05). **(a)** Links significant without the sample covariate only (Lost), in both models (Kept), or with the sample covariate only (Gained). **(b)** Kept and lost links supported by fine-mapped NEPTUNE TUBE eSNPs. **(c)** Log odds that a pair is eQTL-supported by link category (Non-linked, significant in neither model), adjusted for distance to the TSS, sequencing depth, and peak width. **(d)** Change in the log odds that a candidate pair is a significant link when the covariate is added, by the peak's allele-specific chromatin accessibility (ASCA) tier, adjusted for allelic coverage and sequencing depth. **(e)** For peaks that lost a link, ASCA log2(ALT/REF) within heterozygous samples against the genotype effect on accessibility across 22 samples. Lines show least-squares fits, ρ indicates Spearman correlation, and n indicates the number of peaks. **(f)** Log odds that a lost link is eQTL-supported by ASCA tier, adjusted as in (c). In (c), (d), and (f), dots and whiskers show model estimates with 95% CI, dashed lines indicate the reference category (Non-linked in c, and Non-ASCA in d and f), and asterisks indicate P versus the reference (* P < 0.05; ** P < 0.01; *** P < 0.001; ns, not significant).

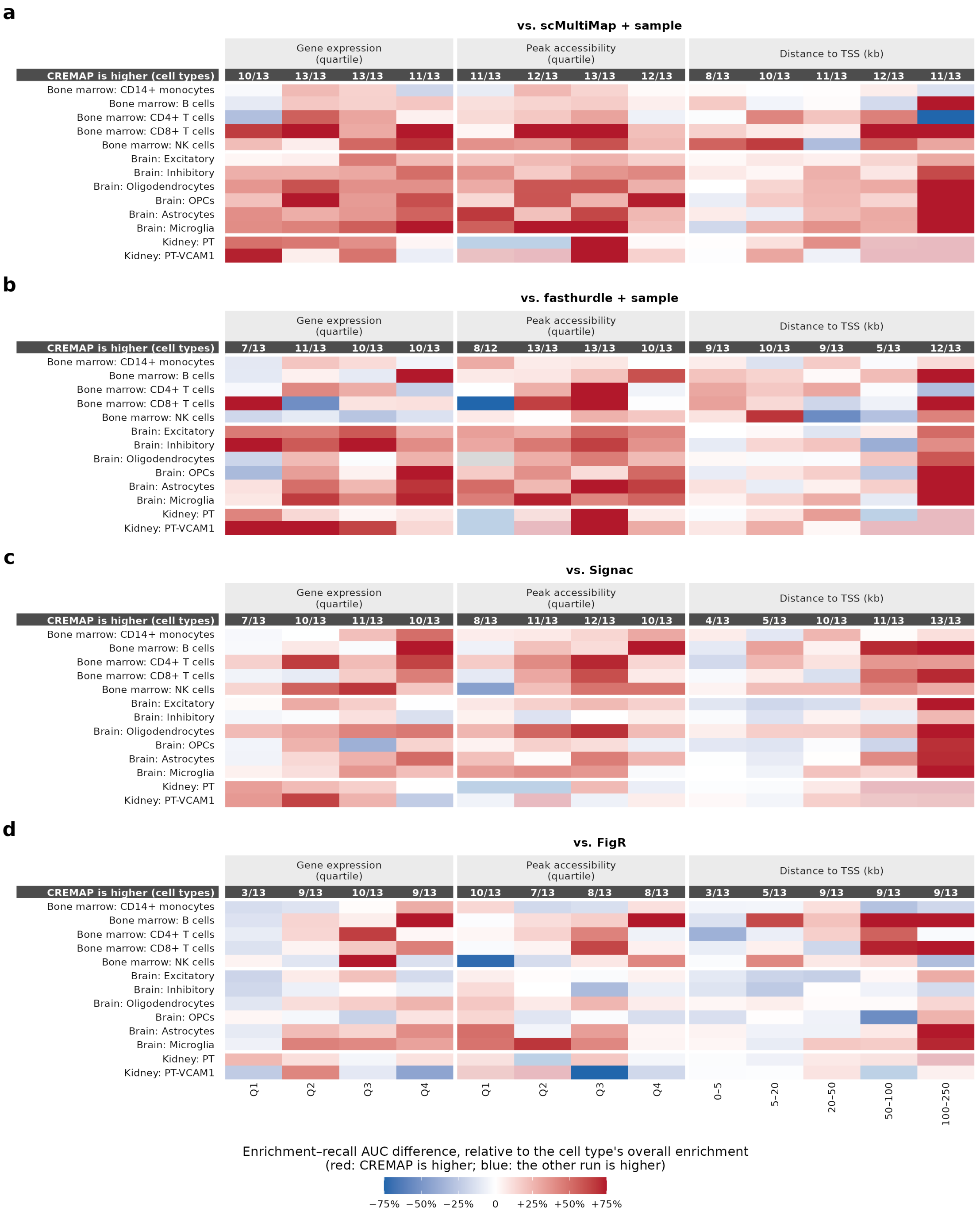


**Figure S7. Performance of CREMAP compared with other methods on multi-sample datasets across different contexts. (a)–(d)** Differences in enrichment–recall AUC between CREMAP and each competing method across candidate-pair strata (columns) for 13 cell types (rows), scaled to each cell type's overall enrichment. Faded cells indicate <100 eQTL-supported pairs in the stratum, and gray cells indicate too few pairs to score. The top row shows the number of scored cell types where CREMAP's AUC is higher.

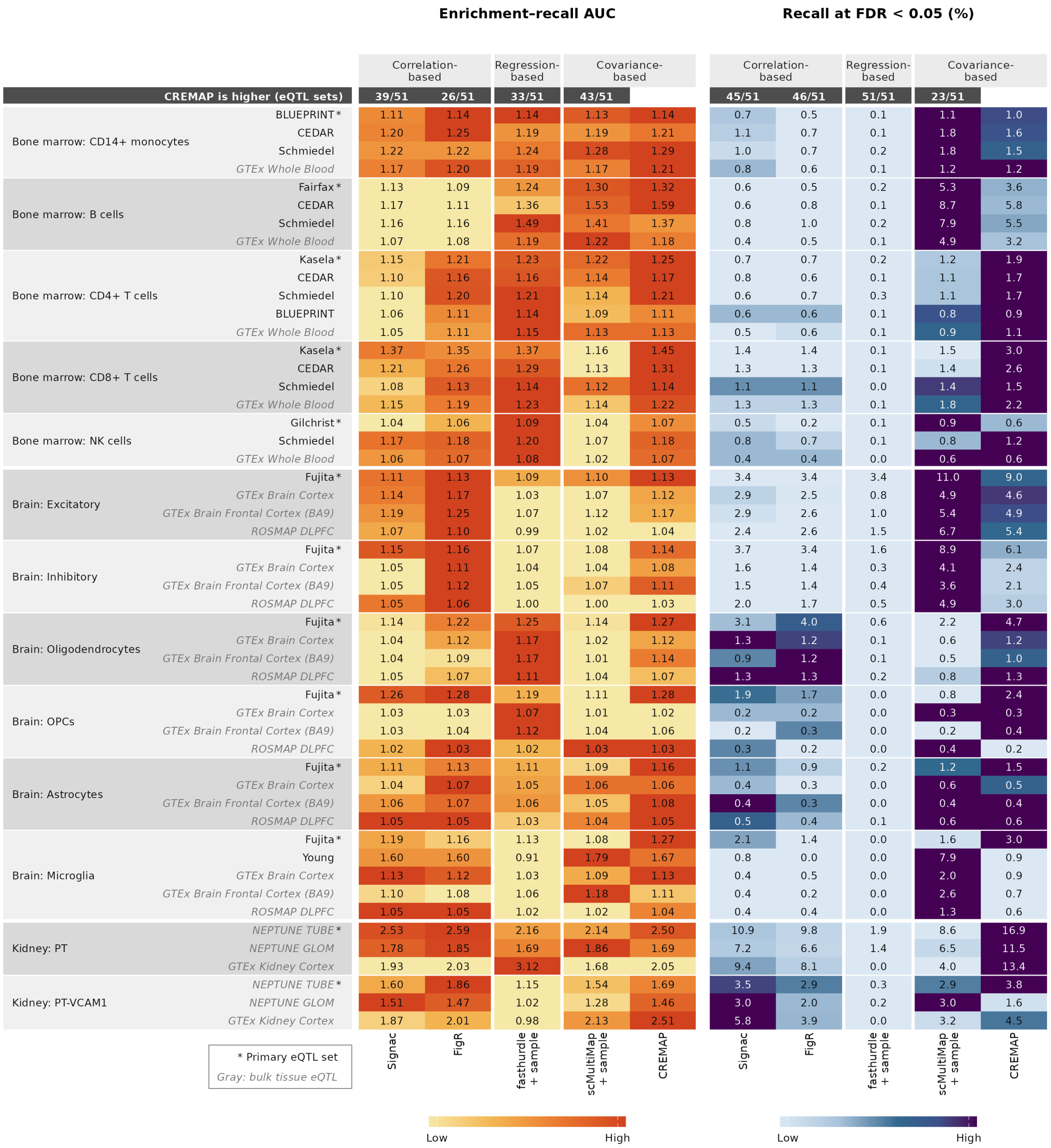


**Figure S8. CREMAP versus other methods on all 51 cell-type-matched eQTL sets.** Enrichment–recall AUC (left) and recall at FDR < 0.05 (right) across five methods for each eQTL set (rows). Color is scaled to the best method in each row, and the top row shows the number of eQTL sets where CREMAP achieves a higher value.

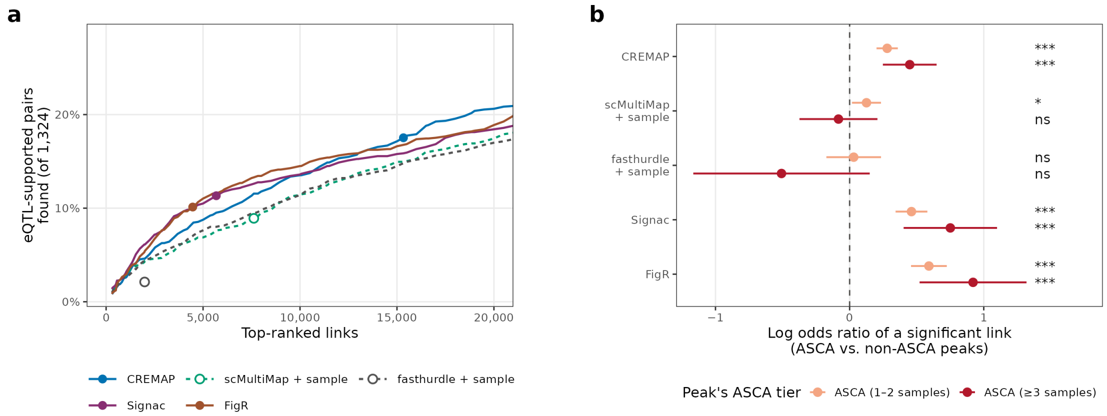


**Figure S9. Comparison of eQTL-supported pairs and links at ASCA peaks across methods.** **(a)** Percentage of eQTL-supported pairs found among the top-ranked links for each method in kidney PT cells. Dots indicate each method's significance call (FDR < 0.05), with fasthurdle called by its two-stage Jiang–Doerge rule (placing its dot below its curve). **(b)** Log odds ratio that a pair is a significant link at ASCA versus non-ASCA peaks (where the vertical dashed line at 0 shows no difference), estimated jointly for the five methods in one model and adjusted for allelic coverage and sequencing depth. Dots and whiskers show estimates with 95% CI, and asterisks denote significance versus non-ASCA peaks (* P < 0.05; ** P < 0.01; *** P < 0.001; ns, not significant).

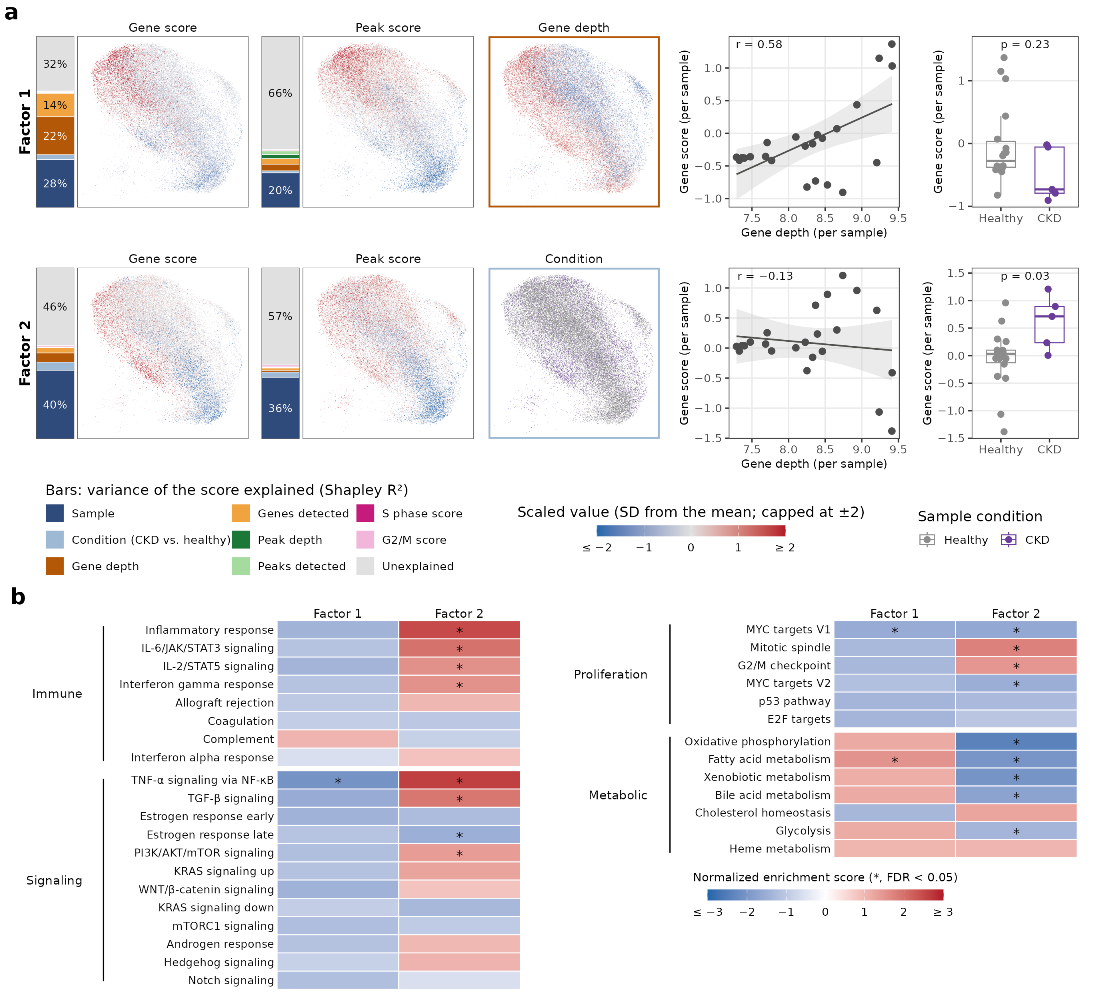

**Figure S10. Technical and biological signals captured by latent factors in kidney proximal tubule cells. (a)** For each factor (rows), panels show the variance of gene and peak scores explained by cell covariates, the scores and strongest covariate after sample (outlined in its legend color) on the WNN UMAP of 35,745 PT cells, and sample-level mean gene scores (23 samples) against mean gene depth and by condition (18 healthy, 5 CKD). In the sample-level plots, the line and shaded band show the least-squares fit with 95% confidence interval, r indicates Pearson correlation, boxes show median and quartiles, and P values are from two-sided Wilcoxon rank-sum tests. **(b)** Normalized enrichment score of selected Hallmark gene sets on each factor's gene loadings.

**Table S1. Links called, recall and enrichment–recall AUC of each method on the validation sets of Fig. 2b.** For each method and validation set (K562 CRISPRi; GM12878 eQTL sets, primary sets marked): links called at FDR < 0.05 among the candidate pairs of the set, positives hit, recall at FDR < 0.05 (%; out of the positives covered by the method's candidate pairs: in K562, 368, and 369 for FigR) and enrichment–recall AUC.

**Table S2. Hallmark gene set enrichment on the gene loadings of the latent factors.** For each cell line, factor (oriented as in Fig. S3) and Hallmark gene set (all 50): the set's process category[61], size, normalized enrichment score and FDR (fgsea, genes preranked by loading; Benjamini–Hochberg within each factor).

**Table S3. Recovery of eQTL-supported pairs at each run's own significance call (kidney PT).** For each run: number of links called (FDR < 0.05; fasthurdle by its two-stage Jiang–Doerge rule), and number and percentage of the eQTL-supported pairs (NEPTUNE TUBE; n = 1,324) and of the eQTL-supported pairs on a peak with significant ASCA (n = 254) among the called links.
